# Sterile protection against *Plasmodium vivax* malaria by repeated blood stage infection in a non-human primate model

**DOI:** 10.1101/2023.02.13.528262

**Authors:** Nicanor Obaldía, Joao Luiz Da Silva Filho, Marlon Núñez, Katherine A. Glass, Tate Oulton, Fiona Achcar, Grennady Wirjanata, Manoj Duraisingh, Philip Felgner, Kevin K.A. Tetteh, Zbynek Bozdech, Thomas D. Otto, Matthias Marti

## Abstract

The malaria parasite *Plasmodium vivax* remains a major global public health challenge, causing major morbidity across tropical and subtropical regions. Several candidate vaccines are in preclinical and clinical trials, however no vaccine against *P. vivax* malaria is approved for use in humans. Here we assessed whether *P. vivax* strain-transcendent immunity can be achieved by repeated infection in *Aotus* monkeys. For this purpose, we repeatedly infected six animals with blood stages of the *P. vivax* Salvador 1 (SAL-1) strain until sterile immune, and then challenged with the AMRU-1 strain. Sterile immunity was achieved in 4/4 *Aotus* monkeys after two homologous infections with the SAL-1 strain, while partial protection against a heterologous AMRU-1 challenge (i.e., delay to infection and reduction in peak parasitemia compared to control) was achieved in 3/3 monkeys. IgG levels based on *P. vivax* lysate ELISA and protein microarray increased with repeated infections and correlated with the level of homologous protection. Analysis of parasite transcriptional profiles across inoculation levels provided no evidence of major antigenic switching upon homologous or heterologous challenge. In contrast, we observed significant transcriptional differences in the *P. vivax* core gene repertoire between SAL-1 and AMRU-1. Together with the strain-specific genetic diversity between SAL-1 and AMRU-1 these data suggest that the partial protection upon heterologous challenge is due to molecular differences between strains (at genome and transcriptome level) rather than immune evasion by antigenic switching. Our study demonstrates that sterile immunity against *P. vivax* can be achieved by repeated homologous blood stage infection in *Aotus* monkeys, thus providing a benchmark to test the efficacy of candidate blood stage *P. vivax* malaria vaccines.

**Author summary:** *Plasmodium vi*vax is the most widespread human malaria parasite. Elimination efforts are complicated due to the peculiar biology of *P. vivax* including dormant liver forms, cryptic reservoirs in bone marrow and spleen and a large asymptomatic infectious reservoir in affected populations. Currently there is no vaccine against malaria caused by *P. viv*ax. Here we induce sterile immunity by repeated *P. vivax* infection with the SAL-1 strain in non-human primates. In contrast, heterologous challenge with the AMRU-1 strain only provided partial protection. Antibody levels against a crude antigen and a protein microarray correlated with the level of homologous protection. Parasite transcriptional profiles across inoculation levels failed to show major antigenic switching across SAL-1 infections or upon heterologous challenge, instead suggesting other mechanisms of immune evasion. Our study demonstrates that sterile immunity against *P. vivax* can be achieved by repeated blood stage infection in *Aotus* monkeys, thus providing a benchmark to test the efficacy of candidate blood stage *P. vivax* malaria vaccines.

## Introduction

Malaria is caused by parasites of the genus *Plasmodium* that are transmitted to humans by the bite of the female anopheles mosquito. Currently, approximately 241 million cases and 0.6 million deaths from malaria occur worldwide, an increase of 12% from the previous year (1). Most deaths are due to infection with *Plasmodium falciparum*, the most pathogenic of the species, especially in children under the age of five living in sub-Saharan Africa (1–3).

After the elimination of *P. falciparum*, *Plasmodium vivax* is expected to remain a major cause of morbidity and mortality outside of Africa, especially in Central and South America, Asia, and the Pacific Islands (4–6). This is due in part to its peculiar biology, including silent parasite liver forms known as hypnozoites that can cause relapses and major parasite reservoirs in bone marrow and spleen that may act as an unobserved pathogenic biomass and source for recrudescence (7–13). Complete removal of the parasite from the human reservoir is therefore challenging (5, 14), underscoring the need for innovative therapeutic strategies including the development of an effective vaccine (15, 16). Vivax malaria impacts the health of individuals of all ages causing repeated febrile episodes and severe anemia (15, 17), clinical severity including hemolytic coagulation disorders, jaundice, coma, acute renal failure, rhabdomyolysis, porphyria, splenic rupture (4, 18), and Acute Respiratory Distress Syndrome (ARDS)(19–22). Fatal *P. vivax* cases are reported from all endemic regions across the globe (3, 15, 23). Compounding the epidemiology of the disease, *P vivax* malaria transmission is intermittent and acquired immunity is short and strain-specific (15). Even in low transmission regions, it is common to find individuals with asymptomatic parasitemia suggestive of natural premunition – a phenomenon resulting from a delicate host-parasite equilibrium in individuals with acquired immunity (15, 24). Epidemiological studies have demonstrated that repeated exposure increases clinical immunity and decreases parasite density and frequency of clinical episodes (25). For instance, individuals subjected to malariotherapy with *P. vivax* for treatment of neurosyphilis rapidly developed immunity after repeated blood stage infections (8, 26–28), and such repeated infection provided strain transcending protection (25). Moreover, acquired immunity by repeated blood stage infection during malariotherapy has been reported in humans against *P. vivax*, *P. falciparum*, *P. ovale* and *P. malariae* (28–30), providing an early benchmark for the feasibility of developing a vaccine against *P. vivax* (8). However, understanding the correlates of protective immunity against *P. vivax* infection has proven difficult, mainly due to the lack of a continuous *in vitro* culture system for this parasite (31, 32).

The development of a vaccine against malaria with at least 75% protective efficacy is one of the two main objectives identified in the roadmap adopted by the global vaccine action plan until 2030 (33). Such an effective *P. vivax* vaccine should provide long-term and strain-transcending immunity. Current *P. vivax* vaccine studies are focused at inducing a stronger antibody response in combination with an already robust T-cell response (34, 35), based upon passive antibody transfer studies done in humans and laboratory animals (25, 36, 37). Nonetheless, to date, there is no vaccine against *P. vivax* approved for use in humans (15).

Several studies suggest that immunity to repeated blood stage infection in non-human primates is strain- and species-specific. For instance, Rhesus macaques immune to one strain of *P. knowlesi* may be partially susceptible to infection by another strain (36). Similar observations have been reported for *Aotus* repeatedly infected with *P. falciparum* blood stage parasites. (38, 39). Interestingly, the same approach has produced heterologous cross-protection against *Plasmodium chabaudi* infection in mice (40). To assess whether strain-transcendent immunity can be achieved by repeated blood stage infection in *P. vivax*, we used the *Aotus* non-human primate model. The aims of our study were to determine i) how many repeated homologous infections are required for control of parasitemia and development of sterile immunity, and ii) whether strain-transcending immunity could be achieved.

## Results

### *P. vivax* blood stage infection induces sterile immunity to homologous challenge

To evaluate the level of protection against repeated *P. vivax* blood stage infection, six *Aotus* monkeys (MN30014, MN30034, MN32028, MN32047, MN25029, MN29012) were infected intravenously with 50,000 parasites of the *P. vivax* SAL-1 strain and monitored until peak parasitemia (**Figure 1A and S1**). SAL-1 strain was originally isolated from a patient in El Salvador in the late 1960s and adapted to *Aotus* monkeys by W.E. Collins (41). During the first infection, all six animals were positive by day 6 post inoculation (PI) and parasitemia increased steadily to more than 100 x 10^3^/µL infected red blood cells (iRBCs)/µL (mean ± sd = 100,198 ± 43,661/µL) until days 13-14 PI, when the animals were treated with a curative course of chloroquine (CQ) (**Figure 1B**). Sixty-five days PI, one animal (MN29012) was removed from the study due to malaria unrelated causes (**Figure 1A and S1**).

**Fig. 1.**
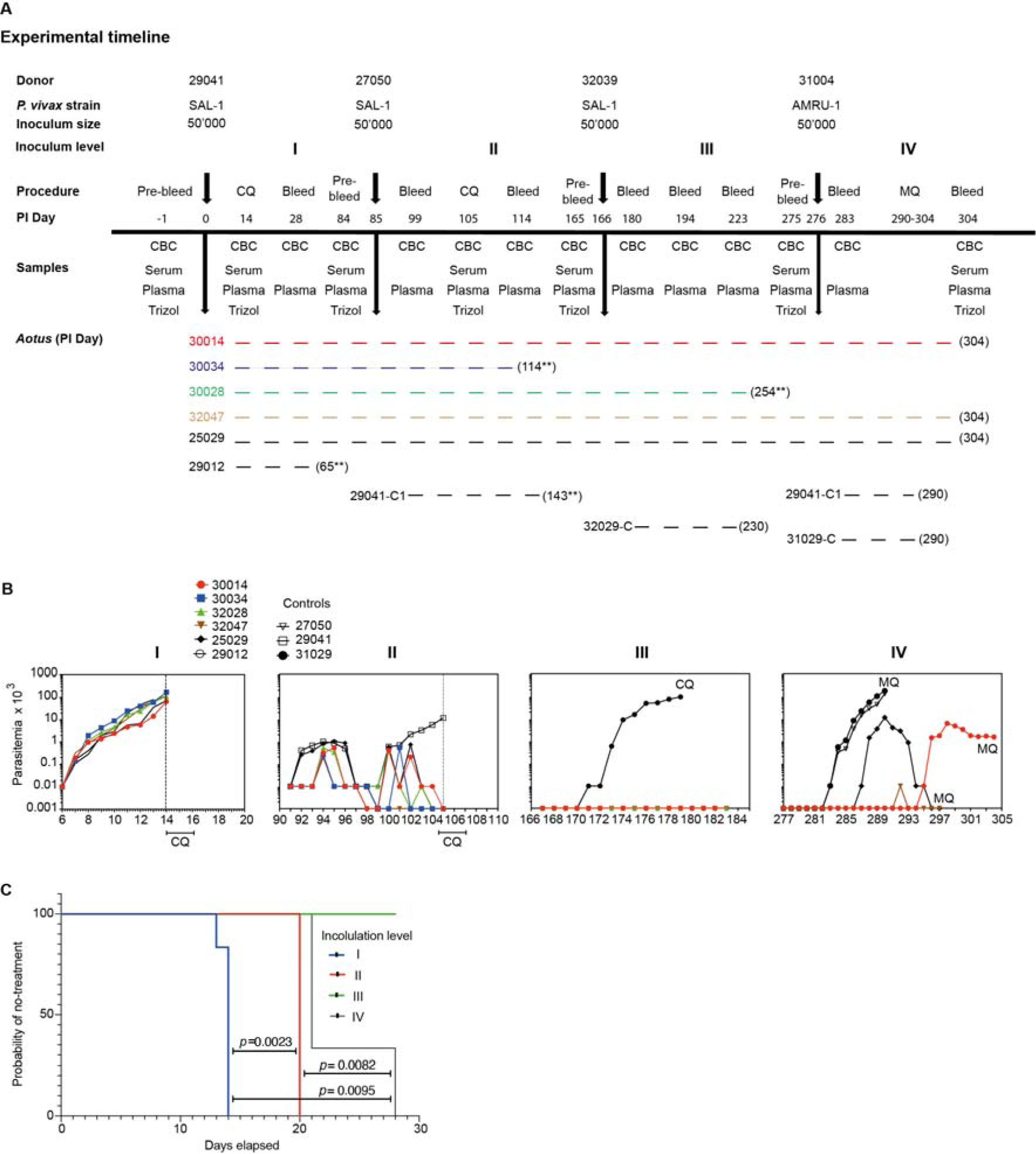
Experimental timeline, parasite dynamics and survival analysis. **A**. Experimental timeline of infection and challenge. *: died of malaria unrelated causes. **: anemia and renal failure. **B**. Peripheral parasitemia across the experiment. Panels I-III show individual parasitemia of *Aotus* monkeys repeatedly infected with *P. vivax* SAL-1 (inoculations I to III). Panel IV shows *Aotus* challenged with *P. vivax* AMRU-1 (inoculation IV). Inoculated control animals were treated at peak parasitemia. **C**. Probability of no treatment of *Aotus* repeatedly infected with the homologous *P. vivax* SAL-1 and heterologous *P. vivax* AMRU-1 strains at each inoculation level. *p* values for survival curve comparison were obtained using the Log-rank (Mantel-Cox) test. Survival curves for homologous infection 1 shown in blue; homologous infection 2 shown in red; homologous infection 3 shown in green. *P. vivax* AMRU-1 heterologous infection 4 shown in black. CBC: red blood cell count. CQ: chloroquine, at 15 mg/kg oral for 3 days. MQ: mefloquine, at 25 mg/kg oral once. C: malaria naïve control. C1: control, once inoculated with *P. vivax*. PI: post inoculation.

Eighty-five days PI the remaining five animals (MN30014, MN30034, MN32028, MN32047, MN25029) and the donor from the first inoculation (MN29041) were infected with SAL-1 using the same inoculum size of 50,000 parasites i.v. This time, by day 91 (day 6 PI of inoculation level II, D6 PI II), all animals were positive by blood smear but parasitemia remained low with a mean peak of 2,332/µL between days 94-95 (D9-10 PI II) (**Figure 1B).** A similar pattern was observed when total parasite load was measured by qPCR (18s rRNA levels) and parasite biomass by ELISA (pLDH levels) after this second inoculation (**Figure S2A, B**). Two animals (MN32047 and MN30034) self-cured on day 98 (D13 PI II) and 102 (D17 PI II) respectively, and a third animal (MN30014) became negative for two days between days 98-99 (D13-14 PI II) but recrudesced on day 100 (D15 PI II) and was treated with CQ on day 105 (D20 PI II) while still positive at the level of <10 parasites /µL. Meanwhile, MN29041 that had controlled its parasitemia until day 98 (D13 PI II), became negative on day 99 (D14 PI II), but recrudesced the next day, reaching a parasitemia level of 11,500/µL on day 105 (D20 PI II) when it was treated with CQ (42). All animals received CQ treatment on day 105 (D20 PI II). Two animals (MN30034 and MN29041) were excluded following CQ treatment - MN30034 on day 169 (D114 PI II) due to severe anemia (Hct% = 20) and kindney failure, and MN29041 for malaria unrelated causes on day 143 (D58 PI II). On day 166 the remaining 4 original animals (MN30014, MN32028, MN32047 and MN25029) plus a malaria naïve infection control (MN32029), were re-inoculated a third time with SAL-1 and followed up as described above (**Figure 1B**). This time, all animals except for the control (MN32029) that had a peak parasitemia of 95,550/µL on day 179 (D13 PI III) remained negative and did not require CQ treatment. Of note, MN32028 had to be removed from the experiment on day 254 (D88 PI III) due to anemia and kidney failure. At necropsy, the animal presented with generalized subcutaneous oedema (Anasarca), with pericardial and pleural effusion, pulmonary oedema and evidence of chronic renal lesions. The cause of death was determined as renal failure (**Figure 1A**).

Altogether, these experiments demonstrate that repeated homologous *P. vivax* infection confers full protection (or sterile immunity) against a homologous challenge.

### Partial protection to heterologous challenge after repeated homologous infection

To determine the difference in protection between homologous and heterologous infections, we challenged on experimental day 276 the three remaining monkeys that went through three SAL-1 inoculations (MN30014, MN32047 and MN25029) plus a new malaria naïve infection control (MN31029) and the donor of the second SAL-1 infection (MN27050) with the CQ resistant AMRU-1 strain (**Figure 1A, B**). The AMRU-I strain was originally isolated from a patient in Papua New Guinea in 1989 (43).

This time all animals became positive. First, the 2 controls (MN27050 and MN31029) were positive on day 283 (D7 PI IV) with peak parasitemia of 131.5 x 10^3^/µL and 180 x 10^3^/µL on day 290 (D14 PI IV), respectively when they were treated with MQ. Meanwhile, MN25029 became positive four days later on day 287 (D11 PI IV) with a lower (10-fold) peak parasitemia of 11.4 x 10^3^/µL on day 290 (D14 PI IV), clearing on day 296 (D20 PI IV) and treated with MQ on day 297 (D21 PI IV). Similarly, MN30014 became positive on day 295 (D19 PI IV) with a 100-fold lower peak parasitemia of 1,700/µL on day 297 (D21 PI IV) compared to the peak parasitemia of the controls. The animal was treated with MQ on day 304 (D28 PI IV) for moderate anemia (Hct% = 27.4) and thrombocytopenia (PLT = 54 x 10^3^/µL), while still positive at 1,510 parasites /µL. In contrast, MN32047 was positive only once on day 292 (D16 PI IV) with less than 10 parasites /µL and was treated with MQ for severe anemia (Hct% = 16) on day 297 (D21 PI IV).

Altogether, this experiment revealed partial protection in 3/3 of the monkeys to a heterologous *P. vivax* challenge in sterile homologous immune animals. Partial protection was characterized by a delay of 4-12 days in patency and reduced parasitemia compared to the controls and a delay of 5-13 days in patency compared to the first homologous SAL-1 challenge. To further investigate the difference between repeated homologous and heterologous infections, we used survival analysis to assess the probability of the test subjects not requiring treatment at each inoculation level (**Figure 1C**). Median time to treatment was established at 14, 20 and none for homologous inoculation levels I-III, respectively, and 21 days for the heterologous challenge. Further analysis of various parasitemia-related parameters including mean days patent, mean day of peak, mean peak parasitemia and the Total Area Under the parasitemia Curve (AUC) (**Figure S2**), indicated that the level of protection against the heterologous challenge in inoculation level IV was similar to protection after one homologous challenge (i.e., inoculation level II). Indeed, the mean days of patency was shorter in infection level IV (unpaired t-test = 3.060; df = 6; *p* = 0.0222) while the mean day to peak parasitemia was longer compared to level II (unpaired t-test = 3.032; df = 6; *p* = 0.0230). No significant difference was found in peak parasitemia (unpaired t-test = 2.191; df = 6; *p* = 0.0709) and AUC (unpaired t-test = 2.409; df = 6; *p* = 0.0526) between level II and IV (**Figure S2**).

### Severe anemia upon *P. vivax* heterologous challenge in sterile homologous immune ***Aotus***

Next, we investigated the longitudinal dynamics of hematological parameters and selected blood chemistry during the repeated *P. vivax* infections (**Figure 2 and Table S1**). During the first inoculation we observed a temporary but significant reduction of both hematocrit and platelet counts that coincided with peak parasitemia, as has been previously observed in *Aotus* (44) and humans experimentally infected with *P. vivax* (45) (**Figure 2A, B - inoculation level I).** During the second homologous infection, and with partial immunity ensuing, all the animals had hematological values within the normal range at peak parasitemia on day 20 PI, when they were treated with CQ for three days (**Figure 2A, B - inoculation level II)**. Of note, MN25029 developed mild anaemia (Hct% = 34.7) and severe thrombocytopenia (39 x 10^3^/µL). During the third homologous infection, none of the animals became parasitemic and their hematocrit and platelet counts remained stable (note MN25029 again developed a moderate thrombocytopenia (90 x 10^3^/µL)) on day 14 PI (**Figure 2A, B – inoculation level III)**. In contrast, the heterologous *P. vivax* AMRU-1 strain challenge triggered anemia and thrombocytopenia in all the animals (**Figure 2A, 2B - inoculation level IV**). For instance, mild to moderate anemia developed in two animals (MN30014 and MN32047) by day 7 PI, even though both animals had undetectable or subpatent parasitemia. Later, on day 28 PI MN30014 developed moderate anemia and severe thrombocytopenia with a parasitemia of 1510/µL and needed treatment with MQ to end the experiment. Similarly, MN32047 developed severe anemia (Hct% = 19.3) on day 18 PI while still negative by light microscopy and needed treatment with MQ on day 21 PI to end the experiment. In contrast, MN25029 developed severe thrombocytopenia (24 x 10^3^/ µL) at peak parasitemia (11,430/µL) on day 14 PI, even though, its Hct% remained within normal limits (Hct% = 45), but later developed moderate anemia (Hct% = 26.2) on day 18 PI when it was still positive at <10 µL and was treated with MQ on day 21 to end the experiment.

**Fig. 2.**
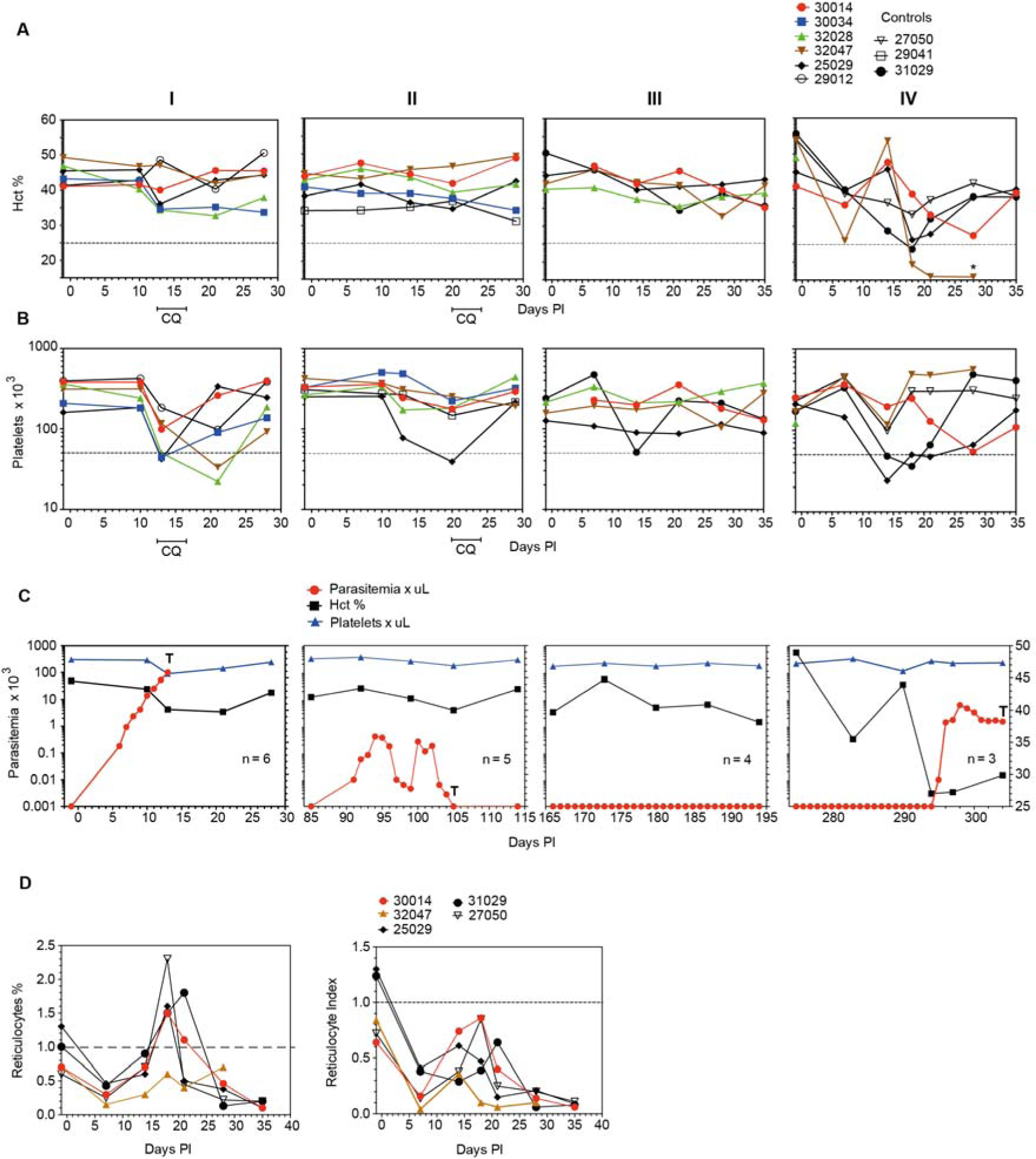
Hematological and parasite parameters. Panels **A-C** show hematocrit levels (Hct%) (**A**), platelet counts (**B**) and combined data from A, B and mean parasitemia (**C**) across inoculation levels I to IV. Panel **D** shows the percentage of reticulocytes and the Reticulocyte Production Index (RPI) at infection level IV. RPI = Reticulocyte Absolute Count/ Reticulocyte Maturation Correction. Reticulocyte Absolute Count = Hct% / 45 x Reticulocyte %. T = CQ: chloroquine, at 15 mg/kg oral for 3 days; MQ at 25 mg/kg once for rescue treatment of *P. vivax* AMRU-1 infections in panel C.

Taken together, these data support previous studies observing the development of severe anemia (hematocrit < 50% of baseline) and thrombocytopenia (< 50 x 10^3^ x µL) in *P. vivax*-infected *Aotus* monkeys around days 12–15 PI (44). Indeed, 2/3 of the remaining original animals (MN30014, MN32047) and a control (inoculated once) (MN27050) showed a Reticulocyte Production Index (RPI) below 1.0 suggestive of bone marrow dyserythropoiesis (46) before inoculation level IV (**Figure 2D**), while only 1/3 of the original animals (MN25029) was over an RPI of 1.0 with a Hct% of 45.

### Antibody levels increase with repeated infections

In a next series of experiments, we analyzed the development of antibodies against a crude *P. vivax* lysate across repeated infections (**Figure 3A, Table S2**). After the first inoculation with *P. vivax* SAL-1 total antibody (Ab) levels reached a mean of 3.1 Log10 arbitrary ELISA units (day 28 PI), decreasing slightly to 2.9 Log10 ELISA units by day 84 PI. After the second homologous inoculation (day 84 PI) Ab levels peaked to 4 Log10 ELISA units on day 114 PI, decreasing slightly again to 3.5 Log10 ELISA units by day 165 PI (**Figure 3B**). After the third homologous inoculation (day 165 PI) when all the animals were sterile protected against challenge (**Figure 3B**), Ab levels remained over 4.0 Log10 ELISA units until day 275 PI when the animals were challenged with the heterologous *P. vivax* AMRU-1 strain. This time a booster response was observed with Ab levels increasing to 4.3 Log10 ELISA units by day 304 PI (**Figure 3B**). Interestingly, Ab levels appear to be negatively correlated with parasitemia (**Figure 3C**). In summary, the dynamics of mean parasitemia and ELISA titers during inoculation levels I-IV suggest that an ELISA titer of 3-4 arbitrary Log10 units would fully protect against challenge with a homologous but only partially protect against a heterologous strain of *P. vivax* (**Figure 3D)**. These correlates of protection provide a benchmark for efficacy testing of *P. vivax* blood stage candidate vaccines in the *Aotus* model.

**Fig. 3.**
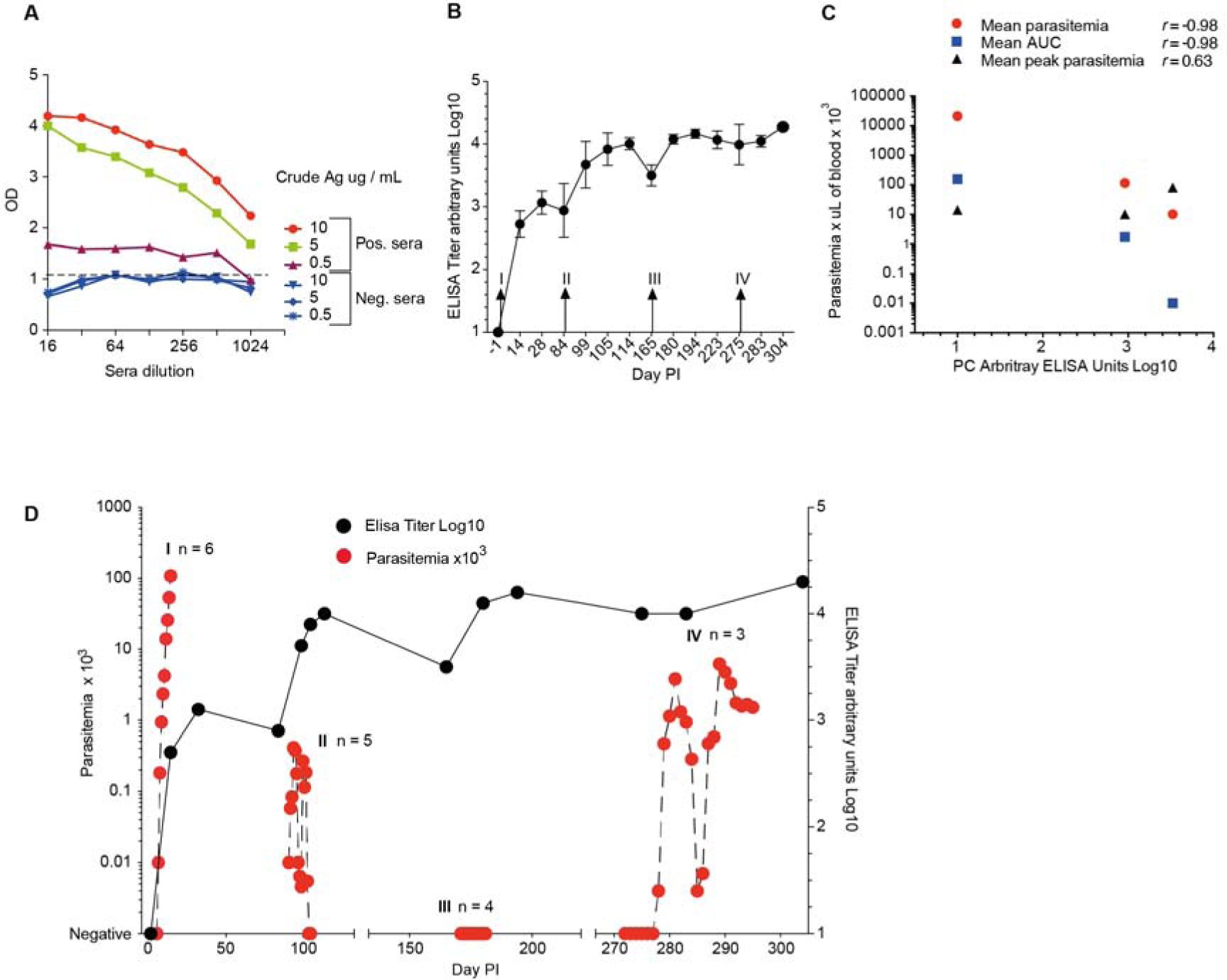
ELISA titers of *Aotus* repeatedly infected with *P. vivax* blood stages. **A.** Crude antigen checkerboard titration. *P. vivax* SAL-1 antigen was prepared from *Aotus* iRBCs purified by Percoll cushion (47%) centrifugation and adsorbed to the plate wells diluted in PBS pH 7.4 at a concentration of 5 µg / mL. Secondary antibodies (peroxidase conjugated Goat anti-monkey, Rhesus macaque) were diluted 1:2000 in PBS pH 7.4., and optical density (OD) read using a 492 nm filter. **B.** Mean ELISA* titers of *Aotus* immunized by repeated infection with the homologous SAL-1 and challenged with the heterologous AMRU-1 strains of *P. vivax*. I-IV indicates inoculation level, each with inoculum of 50 x 10^3^ iRBCs. Level I-III infection with homologous SAL-1. Level IV indicates infection with heterologous AMRU-1. **C.** Pearson correlation analysis of mean ELISA titers at inoculation levels I (n = 6), II (n = 5) and III (n = 4) showed a high negative correlation vs mean parasitemia (*r* – 0.98), the mean area under the curve (AUC) (*r* = −0.98), and a moderate positive correlation vs mean peak parasitemia (*r* = 0.63). **D.** Combined plot of mean parasitemia and ELISA titers with *Aotus* repeatedly infected with the homologous SAL-1 (Infection I-III) and challenged with the heterologous AMRU-1 (Infection IV).

### Quantification of antigen responses using a *P. vivax* protein microarray

Antibody responses during repeated infection (log2 (antigen reactivity/no DNA control reactivity)) show that 66 out of 244 *P. vivax* antigens in the protein microarray demonstrated reactivity above 0 in 10% of all samples analyzed **(Figure 4A).** When we compared the antibody levels for these 66 antigens for all time points in inoculation III (the final homologous challenge) vs inoculation IV (the heterologous challenge) for the three monkeys that completed the entire experiment, there were no differentially reactive antigens (paired t-test with FDR correction). It is possible that there are differentially reactive antigens that were not identified in this study due to the limited number of antigens tested and/or the small sample size.

**Fig. 4.**
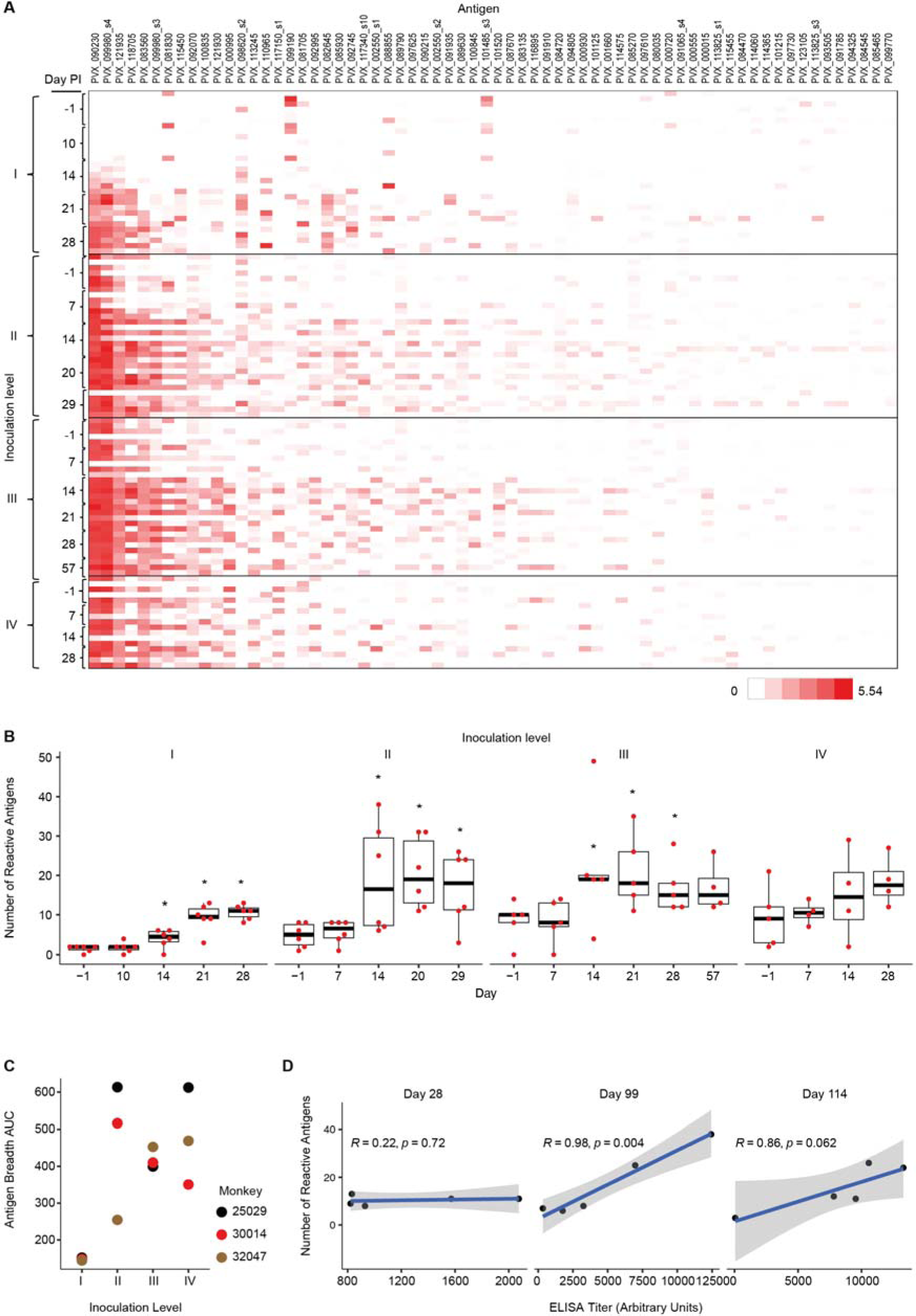
Protein microarray. **A**. Shown are antibody responses (log2(antigen reactivity / no DNA control reactivity)) to 66 out of 244 *P. vivax* IVTT antigens with reactivity above 0 in 10% of all samples and across monkeys. Thus, zero represents equal or lower reactivity than the mean of the no DNA control spots. Antigens are ordered from highest to lowest overall mean. Samples are ordered top to bottom by inoculation level, day, and then by monkey. **B**. Antigen breadth (number of *P. vivax* reactive antigens) by post-infection day at each inoculation level (I-IV). Antigens were considered reactive if the reactivity was higher than the mean + 3SD of the no DNA control spots for that sample. * indicates a significantly higher antigen breadth at that day than at baseline (day −1) within each inoculation (*p*<0.05, Wilcoxon matched pairs test, one-sided). **C.** Area under the curve (AUC) of the antigen breadth at each inoculation level for the three monkeys that completed the experiment**. D.** Pearson correlation of ELISA titer at each day post-infection versus antigen breadth. *p* values shown are from t-tests with the null hypothesis that the correlation coefficient equals 0.

Within inoculation levels I-III, the number of reactive antigens (antigen breadth) was significantly increased at days 14, 21, and/or 28 when compared with the pre-inoculation antigen breadth (**Figure 4B**, *p* < 0.05 Wilcoxon matched pairs test). The trend for increased antigen breadth over time is similar but non-significant for the heterologous infection with the *P. vivax* AMRU-1 in inoculation IV. When we calculated the area under the curve of antigen breadth for each inoculation level for the three monkeys which completed all four inoculations, inoculation III and IV were both significantly higher than inoculation I (and were not different from each other) (**Figure 4C**, *p* < 0.05 repeated measures ANOVA with paired sample post hoc t-tests). These data show that repeated infections of the homologous strain *P. vivax* SAL-1 (inoculation levels I-III) increase the breath of the immune response as the number of infections increased, and that the breadth remained (but did not increase further) high during heterologous challenge with *P. vivax* AMRU-1. Those antigens eliciting the strongest immune response also showed the strongest positive correlation with ELISA titers (**Figure S3**). Interestingly, no negative association with parasite parameters was observed while similar sets of antigens showed significant negative correlations with platelet counts (**Figure S3**). These include two MSP1 peptides (PVX_099980), an ETRAMP peptide (PVX_090230) and peptides to two exported proteins (PVX_121935 and PVX_083560). We also found that the ELISA titer for the crude lysate correlated well with antigen breadth, however correlations were only significant at inoculation level II; days 99 (Pearson R = 0.98, significant at *p* < 0.005) and 114 (Pearson R = 0.86, trend at *p* = 0.062) (**Figure 4D and S4**).

Longitudinal follow up during repeated infection revealed major immunogenic antigens (Ags) by protein microarray. Indeed, seven targets have significantly higher antibody responses at inoculation level III compared to inoculation level I (**Figure S5**, **Table S3**), including the Early Transcribed Membrane Protein (ETRAMP) [PVX_090230], Parasitophorous vacuolar protein 1 (PV1) [PVX_092070], Merozoite Surface Protein 1 (MSP-1) [PVX_099980, fragments 2 & 3], and three Plasmodium Exported Proteins [PVX_121930, PVX_083560 & PVX_121935]. The maintenance of antigen breadth after heterologous challenge (inoculation IV) may suggest the presence of homologous or cross-reactive antigens between the two isolates. However, amino acid sequences for all 7 targets (6 genes) were identical, except for a region of 14 amino acids in one of the Plasmodium exported proteins (PVX_083560). Altogether these data suggest that sterile protection upon homologous challenge and partial protection upon heterologous challenge may not be due to these proteins, however they may be used as correlates of protection.

### Genetic diversity rather than immune evasion determines level of strain transcendent protection

Our data so far suggest that protection from the homologous challenge is antibody mediated, however the limited resolution of the ELISA and protein array data cannot explain the lower protection after the heterologous challenge. As an alternative approach we investigated possible immune evasion mechanisms on genomic and transcriptional level.

For this purpose, we performed whole genome sequencing of both SAL-1 and AMRU-1 strains to improve the strain-transcendent coverage of the existing *P. vivax* microarray platform (47). Selective WGA enabled targeted amplification of the AT-rich subtelomeres of AMRU-1 and SAL-1 strains used in this study (**Figure 5A**). Assembly and annotation generated continuous subtelomere sequences for SAL-1 and AMRU-1. The number of contigs in the original SAL-1 dropped from 2748 to 113, highlighting the continuity of the PacBio assembly (**Figure 5B**). After the annotation with Companion (48) the improved assembly increased the number of *pir* genes for SAL-1 from 124 to 425, demonstrating that long reads better represent the number of variable gene families in subtelomeric regions. Comparison of the *pir* gene repertoire across strains revealed 593 and 425 *pir* genes in AMRU-1 and SAL-1, respectively, compared to over 1000 in the PvP01 reference strain (**Figure 5C**). This difference in number may be because the reference strain came straight from patient infection while SAL-1 and AMRU-1 may have adapted during repeated passages through monkeys. Finally, the proportion of *pir* subtypes remains constant across strains as previously reported (49).

**Fig. 5.**
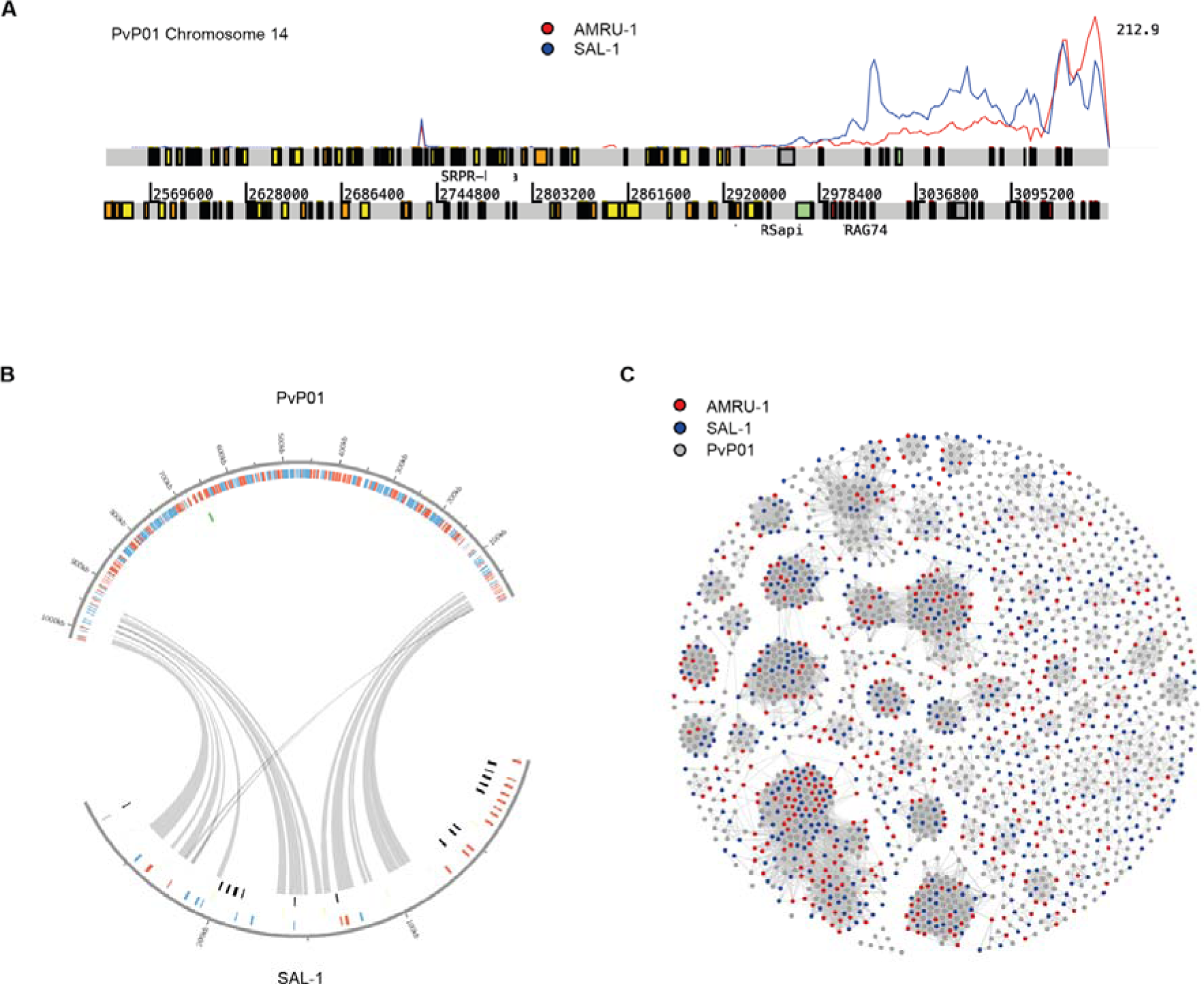
PacBio Whole Genome Sequencing (WGA) of *Plasmodium vivax* SAL-1 and AMRU-1. **A.** Artemis screenshot. Shown is one arm of *P. vivax* PvP01 chromosome 14, with PacBio reads mapped (SAL-1 in blue and AMRU-1 in red). Most of the coverage occurs in subtelomeric regions, demonstrating the specificity of the sWGA. **B**. Circos plot of one representative SAL-1 contig that contains mostly *pir* genes. The contig maps to chromosome 1 of *P. vivax* reference PvP01. Gray lines show syntenic matches of *pir* genes between the two strains. **C.** Gephi plot showing *pir* genes from AMRU-1 (red), SAL-1 (blue) and PvP01 reference (gray). Genes are connected if they share at least 32% global identity.

With this information in hand, we complemented the existing microarray probe set that was generated for the *P. vivax* core genome (47) with probes for the SAL-1 and AMRU-I subtelomeric genes. Next, we investigated whether the virulent phenotype upon heterologous AMRU-1 infection was a result of immune evasion. Differential gene expression (DGE) analysis and principal component analysis (PCA) of the expressed genes revealed greater differences in both core and *pir* genes when comparing AMRU-1 parasites from heterologous challenges (after 3 inoculations with SAL-1) with SAL-1 parasites during the homologous challenges (**Figures 6A – left panel, 6B**). We also compared DGE of AMRU-1 parasites between animals previously infected with three SAL-1 inoculations with i) animals previously infected with only one SAL-1 inoculation and ii) with the malaria naïve infection control. Interestingly, only a small number of changes in core and *pir* genes was observed across these comparisons (**Figure 6A – right panel**). The clear overall similarity of sample distribution in the PCA plots based on DGE of core (**Figure 6B -left panel**) or *pir* genes (**Figure 6B – right panel**) suggests that the repeated SAL-1 infections do not induce extensive *pir* gene switching either in SAL-1 or AMRU-1 parasites. Rather, there appear to be significant strain-specific differences in both core and *pir* expression between SAL-1 and AMRU-1. Further analysis using a *pir* gene network revealed no apparent changes in *pir* gene expression across SAL-1 challenges or upon AMRU-1 challenge (**Figure 6C**).

**Fig. 6.**
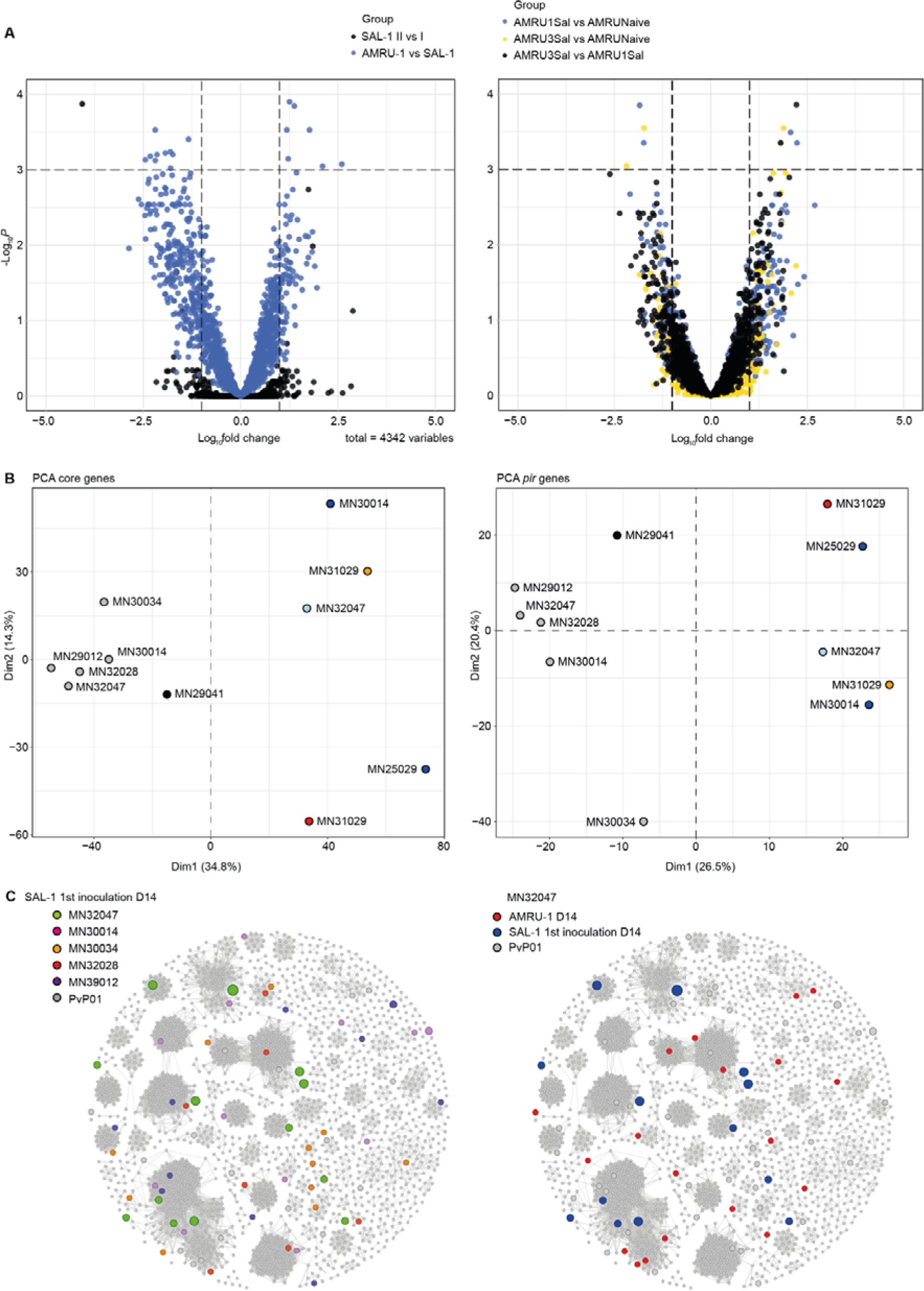
Parasite gene expression comparisons across infection regimes. **A.** Differential gene expression (DGE) across core genes. Volcano plots show DGE between infection regimes. *Left*: DGE of core genes between SAL-1 inoculation II vs SAL-1 inoculation I (black) and between AMRU-1 inoculation IV vs the averaged expression of SAL-1 during the homologous challenges (blue). *Right*: DGE of core genes across AMRU-I infection regimes. Yellow dots represent DGE between AMRU-1 parasites from *Aotus* monkeys previously infected with three SAL-1 inoculations (AMRU3Sal) vs AMRU-1 parasites from naïve *Aotus* monkeys (AMRUNaive). Black dots represent DGE between AMRU3Sal vs AMRU-1 parasites from *Aotus* monkeys previously infected with only one SAL-1 inoculation (AMRU1Sal). Blue dots represent DGE between AMRU1Sal vs AMRUNaive. Each dot represents one annotated *P. vivax* core gene and is displayed according to the fold-change in expression (x-axis, in log2) and statistical significance (y-axis, in negative logarithm to the base 10 of the *p*-value). **B.** Principal Component Analysis (PCA) of the parasite core gene (left panel) and *pir* gene (right panel) expression profiles from each biological replicate, coloured according to the corresponding group: SAL-1 parasites at day 14 PI of the first inoculation (gray); SAL-1 parasites at day 14 PI of the second inoculation (black); AMRU-1 parasites at day 14 PI from *Aotus* monkeys previously infected with three SAL-1 inoculations (light blue dots); gene expression of AMRU-1 parasites at day 28 PI from *Aotus* monkeys previously infected with three SAL-1 inoculations (blue dots); gene expression of AMRU-1 parasites at day 1 PI from naïve *Aotus* monkeys (orange dots); gene expression of AMRU-1 parasites at day 14 PI from naïve *Aotus* monkeys (red dots). **C.** *pir* gene network analysis comparing *P. vivax pir* gene expression in SAL-1 *vs* AMRU-1 infections in *Aotus* monkeys. Same network as in Figure 5C, except that larger circles indicate *pir* gene expression level. Left panel: *pir* expression in SAL-1 parasites at day 14 PI of the first inoculation across individual monkeys. Right panel: comparison of *pir* expression in monkey MN32047 between SAL-1 parasites at day 14 PI of the first inoculation AMRU-1 parasites at day 14 PI of the fourth inoculation.

Altogether, the transcriptional analysis does not indicate that AMRU-1 parasites actively evade the antibody mediated protection induced by SAL-1 homologous challenges by antigenic switching. Thus, the lower protection observed after the heterologous challenge may be due to major genetic and hence antibody epitope variation between these two geographically separated strains (50).

## Discussion

Previous trials of *P. falciparum* and *P. vivax* vaccine candidates have demonstrated the utility of the *Aotus* model in supporting vaccine development (51–53). Various asexual stage vaccine candidate antigens have been subjected to testing in *Aotus* (*52, 54–62*), but only a few have shown some level of efficacy in human clinical trials (15, 63). Development of highly effective strain-transcendent immunity against malaria is a universal goal of vaccine developers (64). Recently, whole organism blood stage malaria vaccines have gained prominence as an alternative to subunit vaccines (65, 66). One major advantage of vaccination using whole blood stage parasites is the multiplicity of immunogenic antigens, including those conserved across strains that may be able to induce strain transcendent immunity (67, 68).

To assess whether strain-transcendent immunity can be achieved by repeated blood stage infection with *P. vivax*, and to investigate possible correlates of protection during repeated infection, we infected six *Aotus* monkeys with the *P. vivax* SAL-1 strain until sterile protected and then challenged with the AMRU-1. We demonstrate that repeated whole blood stage infection with a homologous *P. vivax* strain (i.e., same strain) induces sterile immunity in *Aotus* monkeys after only two infections. In contrast, *Aotus* monkeys infected with *P. falciparum* needed between three to four (69) and six to seven (38) repeated infections, respectively, to achieve sterile immunity. This is consistent with previous observations made during malariotherapy in patients with neurosyphilis demonstrating that immunity to *P. falciparum* is acquired more slowly than to *P. vivax* or *P. malariae* (25). Interestingly, *Saimiri sciureus boliviensis* monkeys immunized with irradiated sporozoites of *P. vivax* SAL-1 and challenged four to nine times with homologous viable sporozoites over a period of almost four years showed sterile protection (70). However, all animals remained susceptible when challenged with SAL-1 blood stage parasites, suggesting that humoral immunity is a correlate of protection against repeated blood stage infections.

Furthermore, our study demonstrates that the sterile immunity achieved after repeated infection with a homologous strain was only partially protective after a heterologous challenge (i.e., delay to infection and reduction in peak parasitemia compared to control). Similar observations have been reported for *P. falciparum* in *Aotus* (38). In both cases, heterologous challenges resulted in severe anemia and thrombocytopenia irrespective of parasitemia level. Such hematological manifestations in semi-immune *Aotus* monkeys with low or subpatent *P. falciparum* parasitemia have been attributed in the past to clearance of non-infected RBCs mediated by autoantibodies (71–74), sequestration of infected RBCs, bone marrow suppression (71, 72) and immune-mediated thrombocytopenia (44, 75–77). The pernicious severe anemia without thrombocytopenia observed in the monkey MN32047 during subpatent parasitemia may have been the result of immune complex disease, or of dyserythropoiesis due to bone marrow infection, as previously described in humans and *Aotus* monkeys infected with *P. falciparum*, *P. malariae and P. brasilianum* (75, 78–84).

A *P. vivax* IVTT protein microarray revealed antibody responses against major immunogenic blood stage antigens (Ags) in this study. Immune reactivity to individual antigens and antibody breath in sera from these animals increased with each inoculation level and were statistically significantly different between inoculation levels I and III, when the animals achieved sterile immunity to a homologous SAL-1 challenge. Among the most significant asexual blood stages antigens detected by the protein microarray were ETRAMP (PVX_090230) located in chromosome 5, and two MSP1 fragments: PVX_099980_s4 and PVX_099980_s2 located on chromosome 7, the latter, a leading vaccine candidate that has been identified as a major determinant of strain-specific protective immunity (85).

In this study, animals with sterile immunity to a *P. vivax* SAL-1 homologous challenge were partially protected against a heterologous AMRU-1 strain. This difference in protection may have been the result of cross-reactive but polymorphic antigens associated with essential parasite phenotypes such as red cell invasion, rosetting or cytoadherence. Maintaining genetic diversity enables immune evasion, as suggested in recent genomic studies of *P. vivax* parasites from distinct geographic origin such as SAL-1 and AMRU-1 (50, 86). Finding conserved and cryptic (not exposed to the immune system) epitopes involved in essential phenotypes that could be targeted by strain transcending neutralizing antibodies represents a possible way forward (87). In contrast to *P. falciparum* that utilizes the variant PfEMP1 antigens to induce cytoadherence and avoid splenic clearance of blood stage parasites, limited vascular sequestration occurs in most other *Plasmodium* species investigated so far. At least in *P. vivax*, this process may be mediated by *P. vivax* orthologs of the *Plasmodium* interspersed repeat (PIR) variant antigens (49). In our study, gene expression analysis along multiple infections allowed correlating *pir* gene expression with the immune response across infections to illuminate parasite immune evasion mechanisms during heterologous challenge. Interestingly, only minor changes in *pir* gene variant expression were observed across all the different inoculation levels, whether homologous or heterologous. Further analysis using a *pir* gene network confirmed no apparent changes in *pir* gene expression in AMRU-1 parasites, regardless of the nature of the previous infections. Together, the transcriptional analysis does not indicate that *P. vivax* actively evades the antibody-mediated protection through antigenic switching. These findings are in accordance with previous studies that have shown no significant difference when comparing the sera of single *versus* repeated infection in patients for VIR antigens and question their role in immune evasion (88, 89). The partial protection observed in the heterologous AMRU-1 challenges may therefore be due to major genetic differences and hence antibody epitope variation between the two strains (50). To overcome this limitation and induce high levels of protective antibodies, we propose use of an immunization regime with whole parasite antigen pools from a mixture of genetically diverse strains.

In conclusion, our study demonstrates that sterile immunity against *P. vivax* can be achieved by repeated homologous blood stage infection in *Aotus* monkeys. It also contributes to our understanding of the pathogenesis of *P. vivax*-induced anemia, *P. vivax* asexual blood stage antigen discovery and correlates of protection, as well as possible immune evasion mechanisms. Most importantly, we establish a benchmark for *P. vivax* protective immunity in the *Aotus* monkey model, providing an important criterion for vaccine development (38).

## Materials and methods

### Ethics statement

The experimental protocol entitled “Induction of sterile protection by blood stage repeated infections in *Aotus* monkeys against subsequent challenge with homologous and heterologous *Plasmodium vivax* strains” was approved and registered at the ICGES Institutional Animal Care and Use Committee (CIUCAL) under accession number CIUCAL-01/2016. The experiment was conducted in accordance with the Animal Welfare Act and the Guide for the Care and Use of Laboratory Animals of the Institute of Laboratory Animal Resources, National Research Council (90), and the laws and regulations of the Republic of Panama.

### Animals and parasites

Twelve laboratory bred (lab-bred) adult male and female “spleen-intact” *Aotus l. lemurinus lemurinus* Panamanian owl monkeys of karyotypes VIII and IX were used in the study (91). The animals were cared and maintained as described elsewhere (92). Isolates of *P. vivax* SAL-1 originally adapted to splenectomized *Aotus* monkeys by W.C. Collins in 1972 (41) and further adapted to spleen intact *A. l. lemurinus* (44, 52), and of *P. vivax* AMRU-1 from Papua New Guinea (PNG) originally adapted to splenectomized *Aotus* by R.D. Cooper in 1994 (93, 94), and further adapted to spleen intact *A. l. lemurinus* by Obaldia N.III. in 1997 (95) were used. This study can be considered as exploratory (i.e. looking for patterns of response rather than hypothesis testing (96)), hence the number of subjects used in the only group studied is typical of such exploratory research with humans (35, 97) and NHP (38).

Briefly, each frozen stabilate of SAL-1 and AMRU-1 was thawed, washed three times with incomplete RPMI medium, and resuspended in 1 ml of RPMI medium. This suspension was used for intravenous (i.v.) inoculation into the saphenous vein of a donor animal using a 25-gauge butterfly needle catheter attached to a 3-ml syringe. When the level of parasitemia reached a peak around days 12-15 post-inoculation (PI), a dilution of blood was made in RPMI to get a total inoculum of 50,000 parasites/ml. All animals received 1 ml of the inoculum through the saphenous vein.

In total 12 spleen intact lab-bred animals were used in this experiment. Six monkeys (three male and three females; MN30014, MN30034, MN32028, MN32047, MN25029, MN29012) were repeatedly infected with *P. vivax* SAL-1 (Homologous challenge) for three times (Levels I-III) and another six animals served as either donors or were assigned as infection or naïve controls. Donors and controls were reassigned back into subsequent inoculation levels as depicted in **Figures 1A** and **S1**. Three SAL-1 homologous sterile immune monkeys from the original six, plus one infected once with SAL-1 and one malaria naive control, were re-challenged (Level IV) with the heterologous CQ resistant and *Aotus* adapted *P. vivax* AMRU-1 strain (44). The animals were treated with CQ at 15 mg/kg for three consecutive days during inoculation level I-III and a drug wash out period of 70 days was kept between inoculation levels I and II and 65 days between levels II and III. No CQ treatment was instituted in inoculation level III. To treat the *P. vivax* AMRU-1 CQ resistant strain, inoculated animals on inoculation level IV and at the end of the experiment were treated with MQ at 25 mg/kg orally once.

### General procedures

Five days after infection, the animals were monitored for any signs of clinical disease and bled 5 µL from a prick made with a lancet in the marginal ear vein to measure daily parasitemia. Parasitemia was determined using thick blood smear stained with Giemsa as described in the Earle and Perez (1932) technique (98). Blood samples were also collected at regular intervals from the femoral vein to assess humoral immune responses against *P. vivax* blood stage proteins, for complete blood count (CBC) and blood chemistry (liver and renal panel), for collection of parasite DNA on FTA^®^ Elute cards (Whatman, Florham Park, NJ. USA) and for RNA in TRizol^®^ solution (Invitrogen, Carlsbad, CA, USA) for molecular biology studies. The animals were treated with Mefloquine (MQ) at 25 mg/kg orally by gastric intubation to end the experiment.

### Criteria for parasitemia

For this study patency was defined as the first of three consecutive positive days after inoculation. Clearance was defined as the first of three consecutive negative days. Recrudescence was defined as the first of three consecutive positive days after a period of clearance. Positivity of <10/µL for less than three days was considered evidence of subpatent infection.

### Criteria for anemia and thrombocytopenia

For this study we classified anemia based on the hematocrit % as mild (Hct% = 31-36), moderate (Hct% = 25-30), or severe (Hct% < 25). Thrombocytopenia was considered mild if platelet counts were between 149-100 x 10^3^/µL, moderate if between 99 and 50 x 10^3^/µL or severe if < 50 x 10^3^/µL.

### Drug treatment

CQ was administered orally for three consecutive days at 10 mg/Kg daily at peak parasitemia. Rescue treatment with MQ was triggered if the hematocrit reached 50% of baseline or hemoglobin was < 8 gm/dL, platelets were < 50 x 10^3^/µL or the animals remained positive by LM after day 28 PI (44).

### Serology

#### Serum ELISA

*P. vivax* SAL-1 antigen was prepared from *Aotus* iRBCs purified by Percoll™(GE Healthcare Bio-Sciences AB, Uppsala, Sweden) cushion (47%) centrifugation as described (99) and adsorbed at 5 μg/mL concentration diluted in PBS pH 7.4 to a 96 well plate at 4-8 degrees Celsius overnight. The plates were blocked with 5% skimmed milk in PBS-0.05% Tween for 2 hours. Serum samples were added to the plate at a dilution of 1/100 in dilution buffer and incubated for one hour, washed further 5 times with PBS pH 7.4 and incubated for one hour with Goat anti-monkey (Rhesus macaque) (Abcam cat # a112767), diluted 1:2000 in PBS pH 7.4. 100 uL per well of the OPD substrate solution (P9029-50G, Sigma-Aldrich, St. Louis, MI, USA) was added to the plate and incubated for 30 minutes away from light and the reaction was stopped with 50 µL of sulfuric acid 3N. To detect the antigen–antibody reactivity, the plates were then read immediately at 492 nm in a ELx808 Plate reader (BioTek^®^, Winooski, VT, USA).

#### pLDH ELISA

To measure *P. vivax* lactate dehydrogenase levels (PvLDH) in the monkey plasma samples, ELISA was performed using a matching pair of capture and detection antibodies (Mybiosource, San Diego, CA). Briefly, 96-well microtiter plate was coated with mouse monoclonal anti-*Plasmodium* LDH (clone #M77288) at a concentration of 2μg/mL in PBS (pH 7.4) and incubated overnight at 4 °C. The plate was washed and incubated with blocking buffer (PBS-BSA 1% - reagent diluent) at room temperature for 2hrs. After washing, samples were diluted 1:2, added to the plate and incubated for 2hrs. Next, plates were washed and HRP-conjugated anti-pLDH detection antibody (clone #M12299), diluted 1:1000 in blocking buffer, was incubated for 1hr at room temperature Plates were washed and incubated for 15 min with substrate solution (OPD), the reaction was stopped adding sulphuric acid 2.5M. Optical density was determined at 450 nm. Cut-off of positivity was defined by correcting absorbance values generated in the plasma samples from blank values (plate controls). Total protein concentration from *Plasmodium falciparum* schizont extracts was determined and samples were used to perform standard curves ranging from 15.625ng/mL up to 2000ng/mL. Lower absorbance values were in the range of O.D = 0.01–0.02. All positive monkey samples gave O.D. values equal to or higher than 0.05.

#### Protein microarray and hybridization

The construction of the protein microarray was conducted using methods as described elsewhere (100). Briefly, coding sequences were PCR amplified from *P. vivax* SAL-1 genomic DNA and cloned into the PXT7 plasmid using homologous recombinant as complete or overlapping fragments, the resulting plasmids (n=244) were expressed in an *Escherichia.coli* based *in vitro* transcription/translation (IVTT) reactions, and the completed reactions printed onto nitrocellulose-coated microarray slides (Grace Bio-Labs, USA). Serum samples mixed with 1/100 blocking buffer (ArrayIt Corp, USA) supplemented with *E. coli* lysate (Genscript, USA). The diluted serum samples were incubated with the protein arrays overnight at 4^0^C, followed by incubation with a goat anti-human IgG Texas Red secondary antibody (Southern Biotech, USA). The arrays were scanned using a Genepix 4300A scanner (Molecular Devices, UK) at 5µm resolution and a wavelength of 594nm (101).

#### Protein microarray data processing and analysis

Raw median fluorescent intensity was local background corrected using the normexp function (offset = 50, method = “mle”, limma R package). All data was log transformed (base 2) and normalized as a ratio of the signal for each spot to the mean of the no DNA control spot within each sample. The number of antigens that have reactivity above 0 in at least 10% of samples was calculated and included in the heatmap (generated in Microsoft excel). Seropositive antigens for each sample were defined as those with reactivity above the mean of the sample specific No DNA control spots + 3SD. These seropositive antigens were totalled for each sample to determine the antigen breadth (number of reactive antigens). The antigen breadth AUC was calculated using the trapezoid rule after limiting the data to only the same number of time points for all inoculations. Pearson’s correlations were performed for available ELISA titers and antigen breadth. All statistics and plots were done using R unless otherwise specified.

### PacBio Whole Genome Sequencing and analysis

*P. vivax* AMRU-1 and SAL-1 were amplified with selective whole genome amplification (sWGA), using primers specific for the subtelomeres that are enriched for low GC content (102). Amplified DNA was used for PacBio sequencing using a commercial protocol (Genscript). We obtained 373,772 subreads with an N50 (type of median length of the reads) of 13,119 bp for SAL-1 and 325,996 subreads with an N50 of 12,035 bp for AMRU-1. The reads were mapped with BWA MEM for quality control (**Figure 5A**) and then assembled with canu (103) (parameter: genomeSize=32m ErrorRate=0.10 gnuplotTested=true useGrid=0 -pacbio-raw, version January 2018). The assemblies generated 113 and 103 contigs with an N50 of 50k and 41k and the largest contig be 195kb and 140kb for SAL-1 and AMRU-1, respectively. For annotation, the assemblies were loaded into Companion (48), using PvP01 as reference strain (June 2018, Augustus cut-off set to 0.4). The genome and its annotation can be found at http://cellatlas.mvls.gla.ac.uk/Assemblies/.

For the Gephi analysis, we extracted all the genes annotated as *pir* from the two Companion runs, merged them with the *pir* genes of PvP01 and performed an all-against-all BLASTp (-F F, Evalue 1e-6). The results were parsed into the open source software Gephi to produce **Figure 5C**. For graphical representation, a force atlas algorithm was run and then the global identity cut-off was set to 32% and the Fruchterman Reingold algorithm was run.

### Gene expression microarray and gene expression analysis

*pir gene probe development*: Sequences from sWGA, representing mostly the AT-rich subtelomeres and excluding the mitochondrial genes, were used as input for probe design using OligoRankPick (104)(oligo size=60, %GC=40). The oligos that were overlapping with core genome oligos from the existing *P. vivax* microarray (47) were removed (12 for SAL-1 and 6 for AMRU-1). The final list of new oligos contains 929 SAL-1 probes and 701 AMRU-1 probes, amongst which 8 match two SAL-1 genes and 5 ma tch two AMRU-1 genes (Full list as **Table S5**).

#### RNA preparation and microarray hybridization

Cell pellets from the blood samples collected at different time points during SAL-1 or AMRU-1 inoculations were stored in trizol. RNA was extracted and processed to be run in a customized microarray assay detecting both core and subtelomoeric genes. The previously described microarray hybridization protocol was used for this study, with several modifications (105). In brief, 100 ng of cDNA was used for subsequent 10 rounds of amplification to generate aminoallyl-coupled cDNA for the hybridizations as described (105). 17µl (∼ 5 µg) of each Cy-5-labelled (GE Healthcare) cDNA of the sample and an equal amount of Cy-3-labelled (GE Healthcare) cDNA of the reference pool were then hybridized together on customized microarray chip using commercially available hybridization platform (Agilent) for 20 h at 70 °C with rotation at 10 rpm. Microarrays were washed and immediately scanned using Power Scanner (Tecan) at 10 µm resolution and with automated photomultiplier tubes gain adjustments to balance the signal intensities between both channels. The reference pool used for microarray was a mixture of 3D7 parasite strain RNA collected every 6 h during 48 h of the full IDC.

#### Microarray analysis

To quantify microarray data signals, intensities were first corrected using an adaptive background correction using the method “normexp” and offset 50 using the Limma package in R(106). Next, we performed within-array loess normalization followed by quantile-normalization between samples/arrays. Each gene expression was estimated as the average of log2 ratios (Cy5/Cy3) of representative probes, thus intensities or log-ratios could be comparable across arrays. Finally, probes with signal showing median foreground intensity less than 2-fold of the median background intensity at either Cy5 (sample RNA) or Cy3 (reference pool RNA) channel were assigned missing values. Fold changes and standard errors of relative gene expression were estimated by fitting a linear model for each gene, followed by empirical Bayes smoothing to the standard errors. Next, the average log 2-expression level for each gene across all the arrays was calculated using the topTable function of the limma package. In parallel we adjusted *p*-values for multiple testing using the Benjamini and Hochberg’s method to control the false discovery rate. The lists of diLerentially expressed genes (DEGs) for each of the comparisons were extracted by defining a cut-off of adjusted *p*-values < 0.001 and fold change > 1. The log fold change and adjusted *p*-values were graphed in volcano plots, using the EnhancedVolcano package in R. From the lists of DEGs, we matched and highlighted those related to 3 categories: (i) surface proteins related to parasite invasion, determined as syntenic orthologs with the *P. falciparum* exportome (refs); (ii) proteins whose expression is spleen-dependent (ref); and (iii) *P. vivax* IVTT antigens inducing high antibody responses as determined by the protein array.

## Acknowledgements

Funding for this study was provided in part by core funds from the Gorgas Memorial Institute of Health Studies and the Sistema Nacional de Investigación de Panamá, SENACYT-SNI awarded to N.O., Panamá, Panamá. This work was also supported by Wolfson Merit Royal Society Award (to M.M.) and Wellcome Trust Center Award 104111 (to F.A., J.L.S.F., T.D.O. and M.M.).

## Author Contributions

**Conceptualization**: Nicanor Obaldia, Matthias Marti.

**Data curation**: Nicanor Obaldia, Joao Luiz Da Silva Filho, Kevin K.A. Tetteh, Thomas D. Otto, Matthias Marti.

**Formal analysis**: Nicanor Obaldia, Joao Luiz Da Silva Filho, Katherine A. Glass, Tate Oulton, Kevin K.A. Tetteh, Thomas D. Otto, Matthias Marti.

**Funding acquisition**: Nicanor Obaldia, Matthias Marti.

**Investigation**: Nicanor Obaldia, Marlon Nuñez, Joao Luiz Da Silva Filho, Katherine A. Glass, Tate Oulton, Fiona Achcar, Grennady Wirjanata, Manoj Duraisingh, Philip Felgner, Kevin K.A. Tetteh, Zbynek Bozdech, Thomas D. Otto, Matthias Marti.

**Methodology**: Nicanor Obaldia, Matthias Marti.

**Project administration**: Nicanor Obaldia.

**Resources**: Nicanor Obaldia, Philip Felgner, Kevin K.A. Tetteh, Zbynek Bozdech, Thomas D. Otto, Matthias Marti.

**Supervision**: Nicanor Obaldia.

**Validation**: Nicanor Obaldia.

**Visualization**: Nicanor Obaldia, Joao Luiz Da Silva Filho, Kevin K.A. Tetteh, Thomas D. Otto, Matthias Marti.

**Writing – original draft**: Nicanor Obaldia, Matthias Marti.

**Writing – review & editing**: Nicanor Obaldia, Joao Luiz Da Silva Filho, Matthias Marti.

## Supplementary figures and legends

**Fig. S1.**
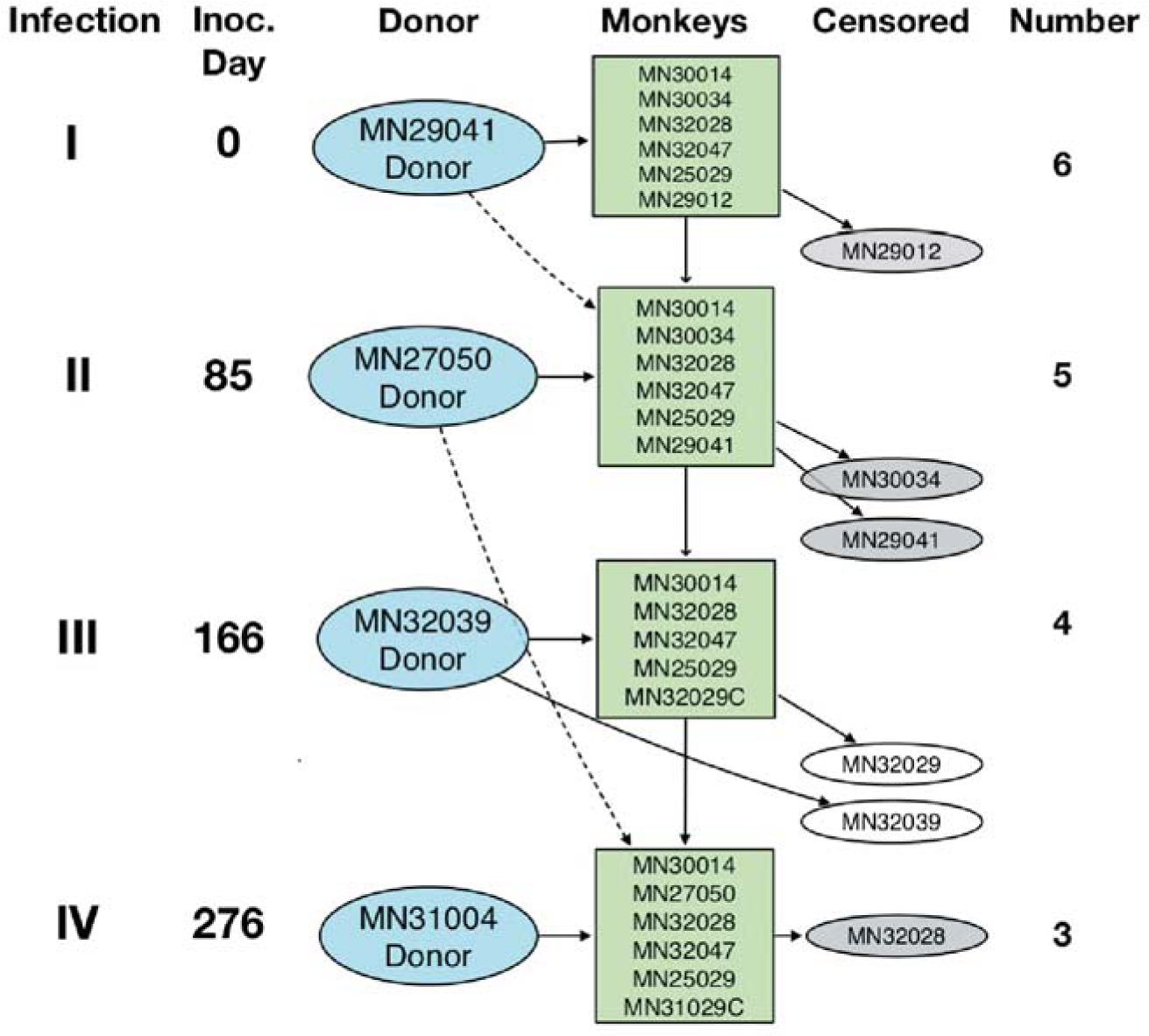
Experimental scheme. Diagram depicting repeated infection of *Aotus* monkeys with the homologous *P. vivax* SAL-1 and challenge with heterologous AMRU-1 strain. Inoculation level, inoculation day, donor monkey, monkey number, and number of animals remaining from the original group of six inoculated are shown.

**Figure S2.**
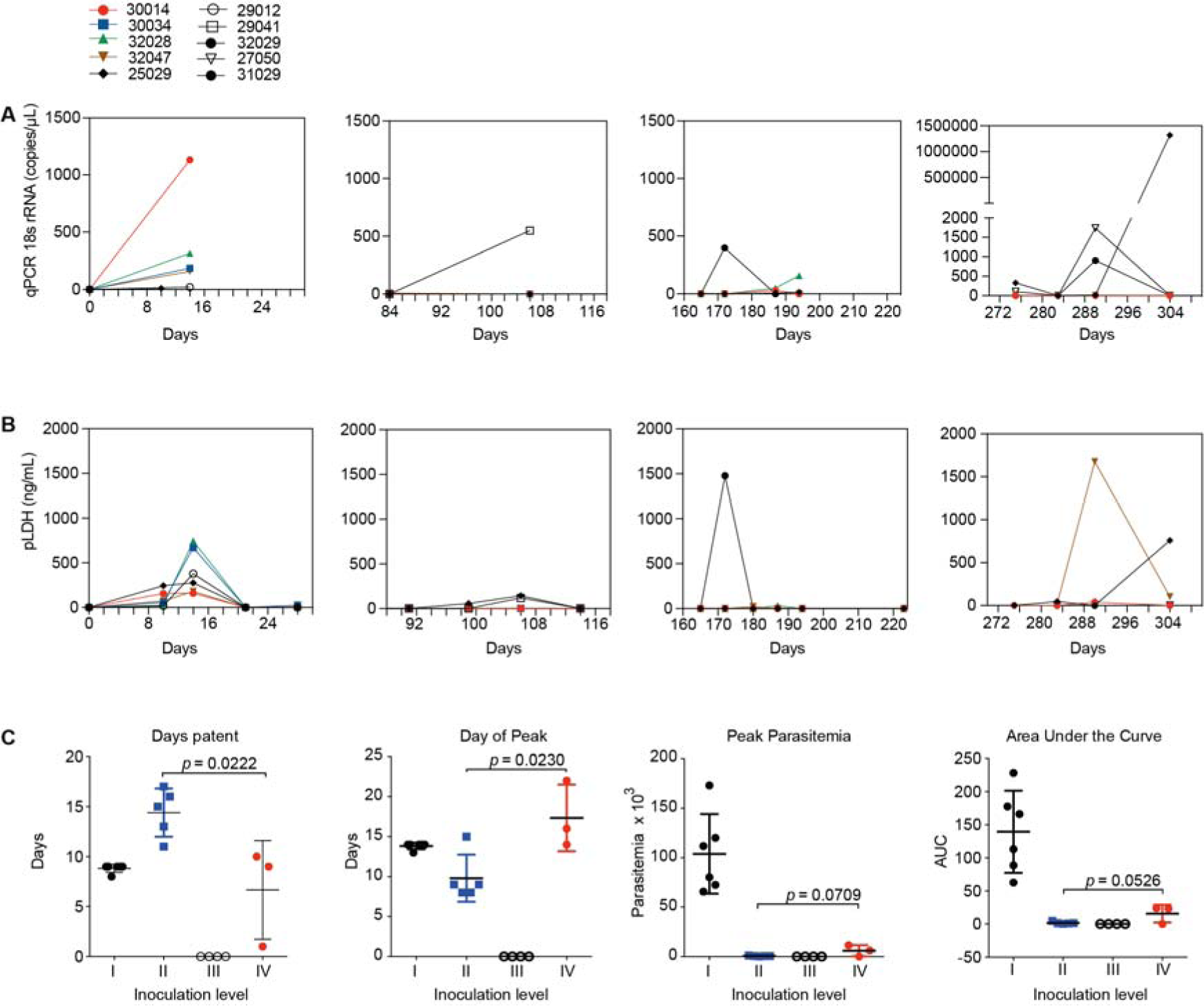
Parasite load and biomass across animals. **A** Parasite load. Panels I-III show parasite load (qPCR 18sRNA in copies x µL) across inoculation levels I-III in individual monkeys infected with *P. vivax* SAL-1. Panel IV shows inoculation level IV, i.e., individual monkeys infected with *P. vivax* AMRU-1. **B**. Parasite biomass. Panels I-III show parasite biomass (pLDH ng/mL) across inoculation levels I-III in individual monkeys infected with *P. vivax* SAL-1. Panel IV shows inoculation level IV, i.e., individual monkeys infected with *P. vivax* AMRU-1. *Plasmodium* LDH levels in ng/mL was calculated based on standard curves using *Plasmodium falciparum* schizont extracts. **C.** Parasitemia parameters across inoculation levels I-IV (Mean ± SD). Left: Days patent. Mid left: Day of Peak. Mid right: Peak parasitemia. Right: Area under the curve (AUC). *P* value; unpaired t-test with equal standard deviation.

**Fig. S3.**
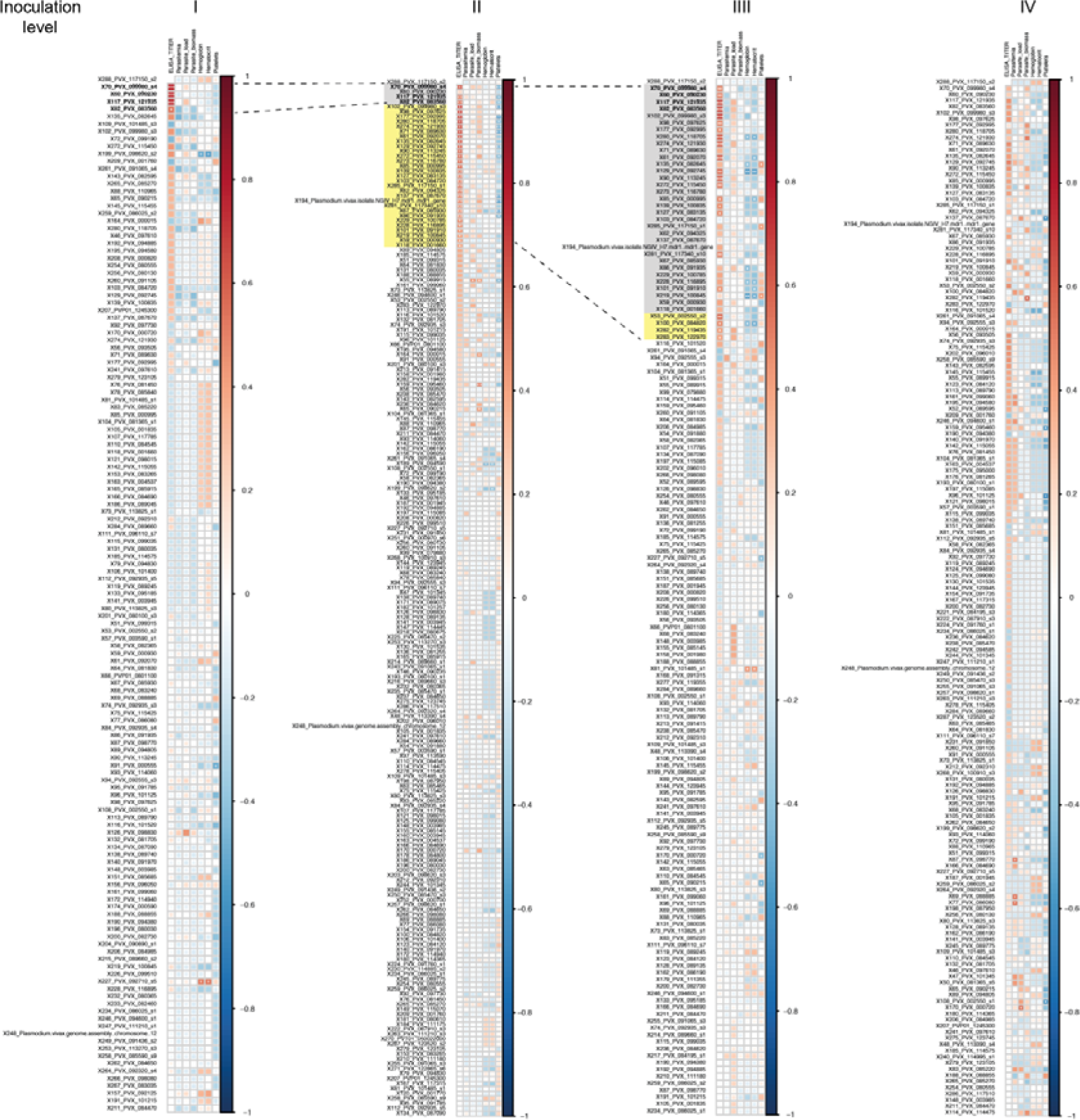
Association of individual antibody responses with ELISA and other parameters. Matrix plot of the Spearman’s rank correlations between the protein array hits and IgG titers (determined by ELISA), parasitemia, parasite load (determined by qPCR), parasite biomass (represented by pLDH levels) and hematological parameters at each inoculation level. Asterisks represent level of significance (*p<0.05, **p<0.01, ***p<0.001).

**Fig. S4.**
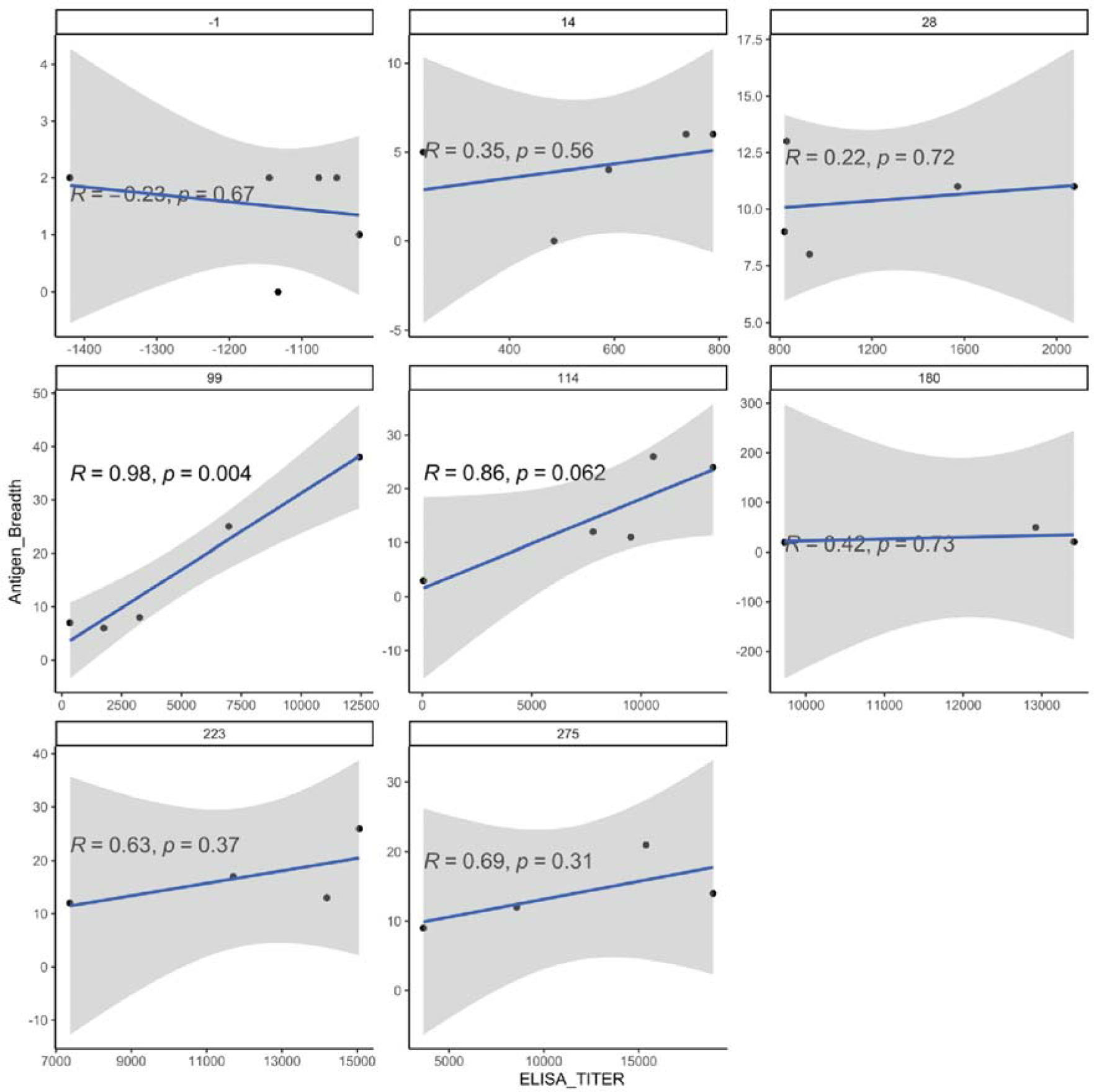
Protein microarray and ELISA titer. Pearson correlation of ELISA titer at each day post-infection vs antigen breadth. *p* values shown are from t-tests with the null hypothesis that the correlation coefficient equals 0.

**Fig. S5.**
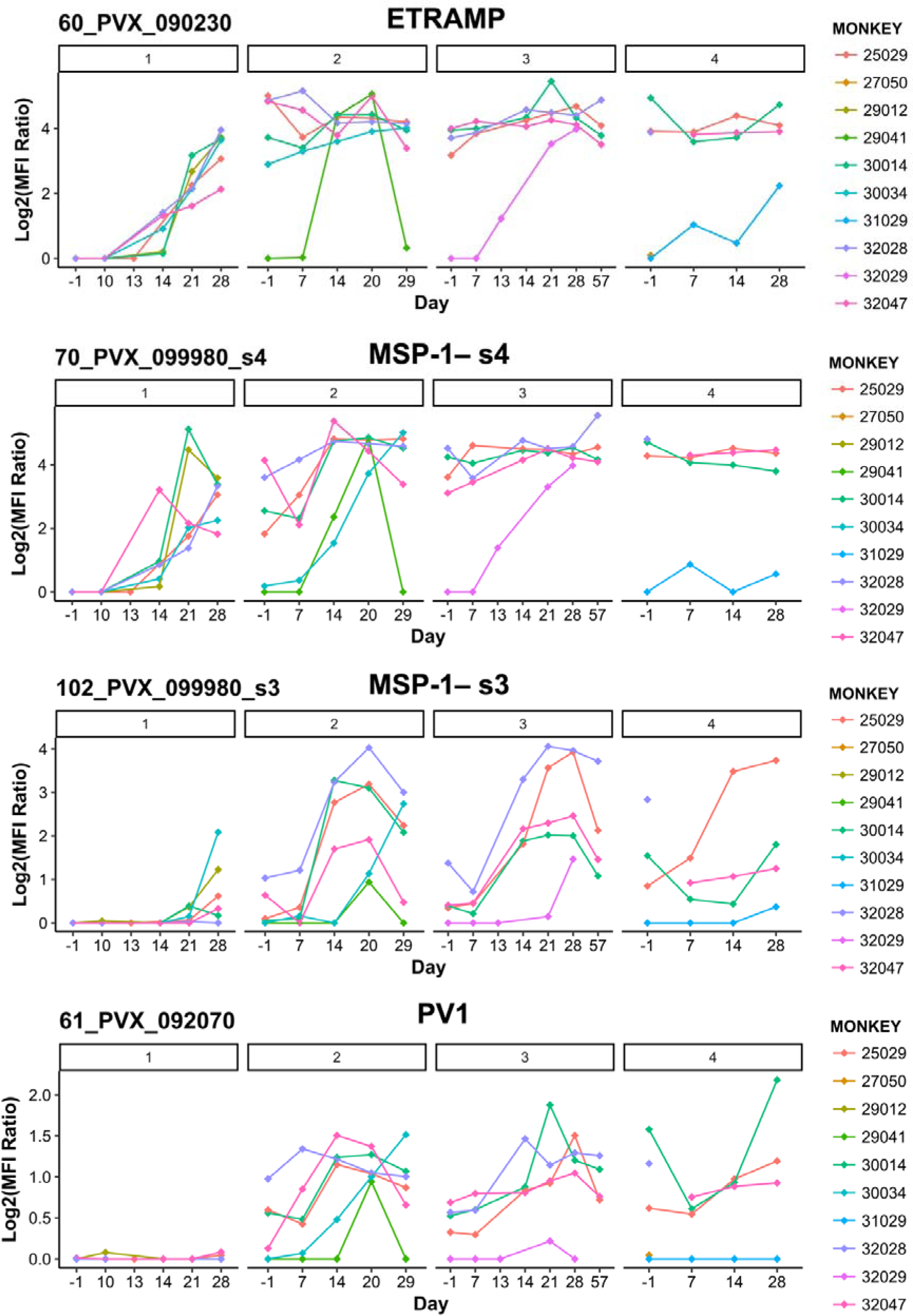
Top protein microarray responses. Reactivity of *Aotus* sera repeatedly infected with *P. vivax* blood stages against selected immunogenic targets. PVX_090230 = ETRAMP; PVX_099980_s3 & PVX_099980_s4 = MSP1; PVX_092070 = PV1. All data is log2(antigen reactivity / no DNA control reactivity). Notably, the dynamic of antibody acquisition varies across individual antigens and monkeys.

## Supplementary tables

**Table S1.** Summary hematological and parasitemia values of *Aotus* repeatedly infected with *P. vivax* SAL-1 (inoculation levels I-III) and challenged with the heterologous *P. vivax* strain AMRU-1 (inoculation level IV).

**Table S2.** Mean ELISA titers of *Aotus* repeatedly infected with *P. vivax* SAL-1 (inoculation levels I-III) and challenged with the heterologous *P. vivax* strain AMRU-1 (inoculation level IV).

**Table S3.** *P. vivax* immunogenic targets with significantly higher antibody levels at inoculation level III vs inoculation level I.

**Table S4:** Normalized microarray data across monkeys and time points.

**Table S5.** Microarray probe set for SAL-1 and AMRU-1.

## References

1. WHO. World Malaria Report 2021. 2021.

2. Schuldt NJ, Amalfitano A. Malaria vaccines: focus on adenovirus based vectors. Vaccine. 2012;30(35):5191–8. Epub 2012/06/12. doi: 10.1016/j.vaccine.2012.05.048. PubMed PMID: 22683663.

3. WHO. World Malaria Report 2019. 2019.

4. Bourgard C, Albrecht L, Kayano A, Sunnerhagen P, Costa FTM. Plasmodium vivax Biology: Insights Provided by Genomics, Transcriptomics and Proteomics. Front Cell Infect Microbiol. 2018;8:34. Epub 20180208. doi: 10.3389/fcimb.2018.00034. PubMed PMID: 29473024; PubMed Central PMCID: PMCPMC5809496.

5. Price RN, Commons RJ, Battle KE, Thriemer K, Mendis K. Plasmodium vivax in the Era of the Shrinking P. falciparum Map. Trends Parasitol. 2020;36(6):560–70.

6. Battle KE, Baird JK. The global burden of Plasmodium vivax malaria is obscure and insidious. PLoS Med. 2021;18(10):e1003799. Epub 2021/10/08. doi: 10.1371/journal.pmed.1003799. PubMed PMID: 34618814; PubMed Central PMCID: PMCPMC8496786.

7. Fernandez-Becerra C, Aparici-Herraiz I, Del Portillo HA. Cryptic erythrocytic infections in Plasmodium vivax, another challenge to its elimination. Parasitol Int. 2021:102527. Epub 2021/12/14. doi: 10.1016/j.parint.2021.102527. PubMed PMID: 34896615.

8. Adams JH, Mueller I. The Biology of Plasmodium vivax. In: Wirth DF, Alonso P, editors. Malaria: Biology in the Era of Eradication. Cold Spring Harbor, NY: Cold Spring Harbor Laboratory Press; 2017. p. 43–54.

9. Bantuchai S, Imad H, Nguitragool W. Plasmodium vivax gametocytes and transmission. Parasitol Int. 2021:102497. Epub 2021/11/09. doi: 10.1016/j.parint.2021.102497. PubMed PMID: 34748969.

10. Baro B, Deroost K, Raiol T, Brito M, Almeida AC, de Menezes-Neto A, et al. Plasmodium vivax gametocytes in the bone marrow of an acute malaria patient and changes in the erythroid miRNA profile. PLoS Negl Trop Dis. 2017;11(4):e0005365. doi: 10.1371/journal.pntd.0005365. PubMed PMID: 28384192; PubMed Central PMCID: PMC5383020.

11. Obaldia N, 3rd, Meibalan E, Sa JM, Ma S, Clark MA, Mejia P, et al. Bone Marrow Is a Major Parasite Reservoir in Plasmodium vivax Infection. MBio. 2018;9(3). doi: 10.1128/mBio.00625-18. PubMed PMID: 29739900; PubMed Central PMCID: PMCPMC5941073.

12. Markus MB. New Evidence for Hypnozoite-Independent Plasmodium vivax Malarial Recurrences. Trends Parasitol. 2018;34(12):1015–6. doi: 10.1016/j.pt.2018.08.010. PubMed PMID: 30213708.

13. Baird JK. African Plasmodium vivax malaria improbably rare or benign. Trends Parasitol. 2022. Epub 2022/06/07. doi: 10.1016/j.pt.2022.05.006. PubMed PMID: 35667992.

14. Baird JK. Resistance to therapies for infection by Plasmodium vivax. Clin Microbiol Rev. 2009;22(3):508–34. Epub 2009/07/15. doi: 10.1128/CMR.00008-09. PubMed PMID: 19597012; PubMed Central PMCID: PMCPMC2708388.

15. De SL, Ntumngia FB, Nicholas J, Adams JH. Progress towards the development of a P. vivax vaccine. Expert Rev Vaccines. 2021. Epub 2021/01/23. doi: 10.1080/14760584.2021.1880898. PubMed PMID: 33481638.

16. White M, Chitnis CE. Potential role of vaccines in elimination of Plasmodium vivax. Parasitol Int. 2022;90:102592. Epub 2022/05/01. doi: 10.1016/j.parint.2022.102592. PubMed PMID: 35489701.

17. White NJ. What causes malaria anemia? Blood. 2022;139(15):2268–9. Epub 2022/04/15. doi: 10.1182/blood.2021015055. PubMed PMID: 35420692.

18. Machado Siqueira A, Lopes Magalhaes BM, Cardoso Melo G, Ferrer M, Castillo P, Martin-Jaular L, et al. Spleen rupture in a case of untreated Plasmodium vivax infection. PLoS Negl Trop Dis. 2012;6(12):e1934. Epub 20121213. doi: 10.1371/journal.pntd.0001934. PubMed PMID: 23272256; PubMed Central PMCID: PMCPMC3521714.

19. Val F, Avalos S, Gomes AA, Zerpa JEA, Fontecha G, Siqueira AM, et al. Are respiratory complications of Plasmodium vivax malaria an underestimated problem? Malar J. 2017;16(1):495. Epub 2017/12/24. doi: 10.1186/s12936-017-2143-y. PubMed PMID: 29273053; PubMed Central PMCID: PMCPMC5741897.

20. Zain Ul A, Qadeer A, Akhtar A, Rasheed A. Severe acute respiratory distress syndrome secondary to Plasmodium vivax malaria. J Pak Med Assoc. 2016;66(3):351–3. Epub 2016/03/13. PubMed PMID: 26968294.

21. Sharma R, Suneja A, Yadav A, Guleria K. Plasmodium vivax Induced Acute Respiratory Distress Syndrome - A Diagnostic and Therapeutic Dilemma in Preeclampsia. J Clin Diagn Res. 2017;11(4):QD03-QD4. Epub 2017/06/03. doi: 10.7860/JCDR/2017/25529.9710. PubMed PMID: 28571216; PubMed Central PMCID: PMCPMC5449862.

22. Lacerda MV, Fragoso SC, Alecrim MG, Alexandre MA, Magalhaes BM, Siqueira AM, et al. Postmortem characterization of patients with clinical diagnosis of Plasmodium vivax malaria: to what extent does this parasite kill? Clin Infect Dis. 2012;55(8):e67–74. Epub 20120706. doi: 10.1093/cid/cis615. PubMed PMID: 22772803.

23. Baird JK. Severe and fatal vivax malaria challenges ‘benign tertian malaria’ dogma. Ann Trop Paediatr. 2009;29(4):251–2. Epub 2009/11/28. doi: 10.1179/027249309X12547917868808. PubMed PMID: 19941746.

24. Mohan K, Stevenson MN. Acquired Immunity to Asexual Blood Stages. In: I.W. S, editor. Malaria: Parasite Biology, Pathogenesis and Protection. Washington D.C.: ASM Press; 1998. p. 467–93.

25. Doolan DL, Dobano C, Baird JK. Acquired immunity to malaria. Clin Microbiol Rev. 2009;22(1):13–36, Table of Contents. Epub 2009/01/13. doi: 10.1128/CMR.00025-08. PubMed PMID: 19136431; PubMed Central PMCID: PMCPMC2620631.

26. Snounou G, Perignon JL. Malariotherapy--insanity at the service of malariology. Adv Parasitol. 2013;81:223–55. Epub 2013/02/07. doi: 10.1016/B978-0-12-407826-0.00006-0. PubMed PMID: 23384625.

27. Mueller I, Galinski MR, Baird JK, Carlton JM, Kochar DK, Alonso PL, et al. Key gaps in the knowledge of Plasmodium vivax, a neglected human malaria parasite. Lancet Infect Dis. 2009;9(9):555–66. Epub 2009/08/22. doi: S1473-3099(09)70177-X [pii] 10.1016/S1473-3099(09)70177-X. PubMed PMID: 19695492.

28. Jeffery GM. Epidemiological significance of repeated infections with homologous and heterologous strains and species of Plasmodium. Bull World Health Organ. 1966;35(6):873–82. Epub 1966/01/01. PubMed PMID: 5298036; PubMed Central PMCID: PMCPMC2476277.

29. Wykes M, Good MF. A case for whole-parasite malaria vaccines. Int J Parasitol. 2007;37(7):705–12. Epub 2007/04/06. doi: 10.1016/j.ijpara.2007.02.007. PubMed PMID: 17408673.

30. Pombo DJ, Lawrence G, Hirunpetcharat C, Rzepczyk C, Bryden M, Cloonan N, et al. Immunity to malaria after administration of ultra-low doses of red cells infected with Plasmodium falciparum. Lancet. 2002;360(9333):610-7. Epub 2002/09/21. doi: 10.1016/S0140-6736(02)09784-2. PubMed PMID: 12241933.

31. Obaldia N, 3rd, Nunez M. On the survival of 48 h Plasmodium vivax Aotus monkey-derived ex vivo cultures: the role of leucocytes filtration and chemically defined lipid concentrate media supplementation. Malar J. 2020;19(1):278. doi: 10.1186/s12936-020-03348-9. PubMed PMID: 32746814.

32. Bermudez M, Moreno-Perez DA, Arevalo-Pinzon G, Curtidor H, Patarroyo MA. Plasmodium vivax in vitro continuous culture: the spoke in the wheel. Malar J. 2018;17(1):301. Epub 20180820. doi: 10.1186/s12936-018-2456-5. PubMed PMID: 30126427; PubMed Central PMCID: PMCPMC6102941.

33. Moorthy VS, Newman RD, Okwo-Bele JM. Malaria vaccine technology roadmap. Lancet. 2013;382(9906):1700-1. Epub 2013/11/19. doi: 10.1016/S0140-6736(13)62238-2. PubMed PMID: 24239252.

34. Hodgson SH, Choudhary P, Elias SC, Milne KH, Rampling TW, Biswas S, et al. Combining Viral Vectored and Protein-in-adjuvant Vaccines Against the Blood-stage Malaria Antigen AMA1: Report on a Phase 1a Clinical Trial. Molecular therapy : the journal of the American Society of Gene Therapy. 2014;22(12):2142–54. doi: 10.1038/mt.2014.157. PubMed PMID: 25156127; PubMed Central PMCID: PMC4250079.

35. Hou MM, Barrett JR, Themistocleous Y, Rawlinson TA, Diouf A, Martinez FJ, et al. Vaccination with Plasmodium vivax Duffy-binding protein inhibits parasite growth during controlled human malaria infection. Sci Transl Med. 2023;15(704):eadf1782. Epub 20230712. doi: 10.1126/scitranslmed.adf1782. PubMed PMID: 37437014.

36. Cohen S, Butcher GA. The immunologic response to plasmodium. Am J Trop Med Hyg. 1972;21(5):713–21. doi: 10.4269/ajtmh.1972.21.713. PubMed PMID: 4561519.

37. Gysin J. Animal Models: Primates. In: Sherman IW, editor. Malaria: Parasite Biology, Pathogenesis and Protection. Washington, D.C.: ASM Press; 1998. p. 419–41.

38. Jones TR, Obaldia N, 3rd, Gramzinski RA, Hoffman SL. Repeated infection of Aotus monkeys with Plasmodium falciparum induces protection against subsequent challenge with homologous and heterologous strains of parasite. Am J Trop Med Hyg. 2000;62(6):675–80. PubMed PMID: 11304053.

39. Siddiqui WA, Taylor DW, Kan SC, Kramer K, Richmond-Crum S. Partial protection of Plasmodium falciparum-vaccinated Aotus trivirgatus against a challenge of a heterologous strain. Am J Trop Med Hyg. 1978;27(6):1277–8. Epub 1978/11/01. doi: 10.4269/ajtmh.1978.27.1277. PubMed PMID: 103450.

40. Elliott SR, Kuns RD, Good MF. Heterologous immunity in the absence of variant-specific antibodies after exposure to subpatent infection with blood-stage malaria. Infect Immun. 2005;73(4):2478–85. Epub 2005/03/24. doi: 10.1128/IAI.73.4.2478-2485.2005. PubMed PMID: 15784594; PubMed Central PMCID: PMCPMC1087398.

41. Collins WE, Contacos PG, Krotoski WA, Howard WA. Transmission of four Central American strains of Plasmodium vivax from monkey to man. J Parasitol. 1972;58(2):332–5. Epub 1972/04/01. PubMed PMID: 4623380.

42. Gomes PS, Bhardwaj J, Rivera-Correa J, Freire-De-Lima CG, Morrot A. Immune Escape Strategies of Malaria Parasites. Front Microbiol. 2016;7:1617. Epub 2016/11/02. doi: 10.3389/fmicb.2016.01617. PubMed PMID: 27799922; PubMed Central PMCID: PMCPMC5066453.

43. Rieckmann KH, Davis DR, Hutton DC. Plasmodium vivax resistance to chloroquine? Lancet. 1989;2(8673):1183-4. Epub 1989/11/18. doi: 10.1016/s0140-6736(89)91792-3. PubMed PMID: 2572903.

44. Obaldia N, 3rd. Clinico-pathological observations on the pathogenesis of severe thrombocytopenia and anemia induced by Plasmodium vivax infections during antimalarial drug efficacy trials in Aotus monkeys. The American journal of tropical medicine and hygiene. 2007;77(1):3–13. Epub 2007/07/11. PubMed PMID: 17620623.

45. Woolley SD, Marquart L, Woodford J, Chalon S, Moehrle JJ, McCarthy JS, et al. Haematological response in experimental human Plasmodium falciparum and Plasmodium vivax malaria. Malar J. 2021;20(1):470. Epub 2021/12/22. doi: 10.1186/s12936-021-04003-7. PubMed PMID: 34930260; PubMed Central PMCID: PMCPMC8685492.

46. Wickramasinghe SN, Looareesuwan S, Nagachinta B, White NJ. Dyserythropoiesis and ineffective erythropoiesis in Plasmodium vivax malaria. Br J Haematol. 1989;72(1):91–9. PubMed PMID: 2660903.

47. Bozdech Z, Mok S, Hu G, Imwong M, Jaidee A, Russell B, et al. The transcriptome of Plasmodium vivax reveals divergence and diversity of transcriptional regulation in malaria parasites. Proc Natl Acad Sci U S A. 2008;105(42):16290–5. Epub 2008/10/15. doi: 0807404105 [pii] 10.1073/pnas.0807404105. PubMed PMID: 18852452; PubMed Central PMCID: PMCPMC2571024.

48. Steinbiss S, Silva-Franco F, Brunk B, Foth B, Hertz-Fowler C, Berriman M, et al. Companion: a web server for annotation and analysis of parasite genomes. Nucleic Acids Res. 2016;44(W1):W29–34. Epub 20160421. doi: 10.1093/nar/gkw292. PubMed PMID: 27105845; PubMed Central PMCID: PMCPMC4987884.

49. Auburn S, Bohme U, Steinbiss S, Trimarsanto H, Hostetler J, Sanders M, et al. A new Plasmodium vivax reference sequence with improved assembly of the subtelomeres reveals an abundance of pir genes. Wellcome Open Res. 2016;1:4. Epub 20161115. doi: 10.12688/wellcomeopenres.9876.1. PubMed PMID: 28008421; PubMed Central PMCID: PMCPMC5172418.

50. Buyon LE, Santamaria AM, Early AM, Quijada M, Barahona I, Lasso J, et al. Population genomics of Plasmodium vivax in Panama to assess the risk of case importation on malaria elimination. PLoS Negl Trop Dis. 2020;14(12):e0008962. Epub 2020/12/15. doi: 10.1371/journal.pntd.0008962. PubMed PMID: 33315861; PubMed Central PMCID: PMCPMC7769613 was unable to confirm their authorship contributions. On their behalf, the corresponding author has reported their contributions to the best of their knowledge.

51. Collins WE. The Owl Monkey as a Model for Malaria. In: Baer JFW, R.E.; Kakoma, I., editor. Aotus The Owl Monkey. San Diego, CA: Academic Press Inc.; 1994. p. 217-44.

52. Obaldia N, 3rd, Stockelman MG, Otero W, Cockrill JA, Ganeshan H, Abot EN, et al. A Plasmodium vivax Plasmid DNA- and Adenovirus-Vectored Malaria Vaccine Encoding Blood-Stage Antigens AMA1 and MSP142 in a Prime/Boost Heterologous Immunization Regimen Partially Protects Aotus Monkeys against Blood-Stage Challenge. Clin Vaccine Immunol. 2017;24(4). doi: 10.1128/CVI.00539-16. PubMed PMID: 28179404; PubMed Central PMCID: PMCPMC5382831.

53. Douglas AD, Baldeviano GC, Lucas CM, Lugo-Roman LA, Crosnier C, Bartholdson SJ, et al. A PfRH5-based vaccine is efficacious against heterologous strain blood-stage Plasmodium falciparum infection in aotus monkeys. Cell Host Microbe. 2015;17(1):130–9. doi: 10.1016/j.chom.2014.11.017. PubMed PMID: 25590760; PubMed Central PMCID: PMCPMC4297294.

54. Gramzinski RA, Millan CL, Obaldia N, Hoffman SL, Davis HL. Immune response to a hepatitis B DNA vaccine in Aotus monkeys: a comparison of vaccine formulation, route, and method of administration. Mol Med. 1998;4(2):109–18. Epub 1998/04/16. PubMed PMID: 9508788; PubMed Central PMCID: PMC2230303.

55. Herrera MA, Rosero F, Herrera S, Caspers P, Rotmann D, Sinigaglia F, et al. Protection against malaria in Aotus monkeys immunized with a recombinant blood-stage antigen fused to a universal T-cell epitope: correlation of serum gamma interferon levels with protection. Infect Immun. 1992;60(1):154–8. PubMed PMID: 1370271; PubMed Central PMCID: PMCPMC257516.

56. Herrera S, Herrera MA, Certa U, Corredor A, Guerrero R. Efficiency of human Plasmodium falciparum malaria vaccine candidates in Aotus lemurinus monkeys. Mem Inst Oswaldo Cruz. 1992;87 Suppl 3:423–8. PubMed PMID: 1343722.

57. Herrera S, Herrera MA, Corredor A, Rosero F, Clavijo C, Guerrero R. Failure of a synthetic vaccine to protect Aotus lemurinus against asexual blood stages of Plasmodium falciparum. Am J Trop Med Hyg. 1992;47(5):682–90. PubMed PMID: 1449209.

58. Inselburg J, Bzik D, Li W, Green K, Kansopon J, Hahm B, et al. Protective immunity induced in Aotus monkeys by recombinant SERA proteins of Plasmodium falciparum. Infect Immun. 1991;59:1247–50.

59. Jones TR, Narum DL, Gozalo AS, Aguiar J, Fuhrmann SR, Liang H, et al. Protection of Aotus monkeys by Plasmodium falciparum EBA-175 region II DNA prime-protein boost immunization regimen. J Infect Dis. 2001;183(2):303–12. doi: 10.1086/317933. PubMed PMID: 11110648.

60. Jones TR, Obaldia N, 3rd, Gramzinski RA, Charoenvit Y, Kolodny N, Kitov S, et al. Synthetic oligodeoxynucleotides containing CpG motifs enhance immunogenicity of a peptide malaria vaccine in Aotus monkeys. Vaccine. 1999;17(23-24):3065–71. Epub 1999/08/26. PubMed PMID: 10462241.

61. Siddiqui WA. An effective immunization of experimental monkeys against a human malaria parasite, Plasmodium falciparum. Science. 1977;197(4301):388-9. Epub 1977/07/22. doi: 10.1126/science.406671. PubMed PMID: 406671.

62. Singh S, Miura K, Zhou H, Muratova O, Keegan B, Miles A, et al. Immunity to recombinant plasmodium falciparum merozoite surface protein 1 (MSP1): protection in Aotus nancymai monkeys strongly correlates with anti-MSP1 antibody titer and in vitro parasite-inhibitory activity. Infect Immun. 2006;74(8):4573–80. Epub 2006/07/25. doi: 10.1128/IAI.01679-05. PubMed PMID: 16861644; PubMed Central PMCID: PMC1539572.

63. Beeson JG, Kurtovic L, Dobano C, Opi DH, Chan JA, Feng G, et al. Challenges and strategies for developing efficacious and long-lasting malaria vaccines. Sci Transl Med. 2019;11(474). Epub 2019/01/11. doi: 10.1126/scitranslmed.aau1458. PubMed PMID: 30626712.

64. Elias SC, Collins KA, Halstead FD, Choudhary P, Bliss CM, Ewer KJ, et al. Assessment of immune interference, antagonism, and diversion following human immunization with biallelic blood-stage malaria viral-vectored vaccines and controlled malaria infection. J Immunol. 2013;190(3):1135–47. Epub 20130104. doi: 10.4049/jimmunol.1201455. PubMed PMID: 23293353; PubMed Central PMCID: PMCPMC3672846.

65. Stanisic DI, Good MF. Whole organism blood stage vaccines against malaria. Vaccine. 2015;33(52):7469–75. Epub 2015/10/03. doi: 10.1016/j.vaccine.2015.09.057. PubMed PMID: 26428451.

66. Good MF, Stanisic DI. Whole parasite vaccines for the asexual blood stages of Plasmodium. Immunol Rev. 2020;293(1):270–82. Epub 2019/11/12. doi: 10.1111/imr.12819. PubMed PMID: 31709558.

67. Raja AI, Cai Y, Reiman JM, Groves P, Chakravarty S, McPhun V, et al. Chemically Attenuated Blood-Stage Plasmodium yoelii Parasites Induce Long-Lived and Strain-Transcending Protection. Infect Immun. 2016;84(8):2274–88. Epub 2016/06/02. doi: 10.1128/IAI.00157-16. PubMed PMID: 27245410; PubMed Central PMCID: PMCPMC4962623.

68. Stanisic DI, Ho MF, Nevagi R, Cooper E, Walton M, Islam MT, et al. Development and Evaluation of a Cryopreserved Whole-Parasite Vaccine in a Rodent Model of Blood-Stage Malaria. mBio. 2021;12(5):e0265721. Epub 2021/10/20. doi: 10.1128/mBio.02657-21. PubMed PMID: 34663097; PubMed Central PMCID: PMCPMC8524336.

69. Udeinya IJ, Miller LH, McGregor IA, Jensen JB. Plasmodium falciparum strain-specific antibody blocks binding of infected erythrocytes to amelanotic melanoma cells. Nature. 1983;303(5916):429-31. doi: 10.1038/303429a0. PubMed PMID: 6343885.

70. Collins WE, Skinner JC, Millet P, Broderson JR, Filipski VK, Morris CL, et al. Reinforcement of immunity in Saimiri monkeys following immunization with irradiated sporozoites of Plasmodium vivax. Am J Trop Med Hyg. 1992;46(3):327–34. doi: 10.4269/ajtmh.1992.46.327. PubMed PMID: 1558272.

71. Egan AF, Fabucci ME, Saul A, Kaslow DC, Miller LH. Aotus New World monkeys: model for studying malaria-induced anemia. Blood. 2002;99(10):3863–6. Epub 2002/05/03. doi: 10.1182/blood.v99.10.3863. PubMed PMID: 11986251.

72. Jones TR, Stroncek DF, Gozalo AS, Obaldia N, 3rd, Andersen EM, Lucas C, et al. Anemia in parasite- and recombinant protein-immunized aotus monkeys infected with Plasmodium falciparum. Am J Trop Med Hyg. 2002;66(6):672–9. Epub 2002/09/13. doi: 10.4269/ajtmh.2002.66.672. PubMed PMID: 12224573.

73. Mourao LC, Cardoso-Oliveira GP, Braga EM. Autoantibodies and Malaria: Where We Stand? Insights Into Pathogenesis and Protection. Front Cell Infect Microbiol. 2020;10:262. Epub 2020/07/01. doi: 10.3389/fcimb.2020.00262. PubMed PMID: 32596165; PubMed Central PMCID: PMCPMC7300196.

74. Mourao LC, Medeiros CMP, Cardoso-Oliveira GP, Roma P, Aboobacar J, Rodrigues BCM, et al. Effects of IgG and IgM autoantibodies on non-infected erythrocytes is related to ABO blood group in Plasmodium vivax malaria and is associated with anemia. Microbes Infect. 2020;22(8):379–83. Epub 2020/02/26. doi: 10.1016/j.micinf.2020.02.003. PubMed PMID: 32097712.

75. Weller RE, Collins WE, Buschbom RL, Malaga CA, Ragan HA. Impaired renal function in owl monkeys (Aotus nancymai) infected with Plasmodium falciparum. Mem Inst Oswaldo Cruz. 1992;87 Suppl 3:435–42. Epub 1992/01/01. doi: 10.1590/s0074-02761992000700073. PubMed PMID: 1343724.

76. Nagatake T, Broderson JR, Tegoshi T, Collins WE, Aikawa M. Renal pathology in owl monkeys vaccinated with Plasmodium falciparum asexual blood-stage synthetic peptide antigens. Am J Trop Med Hyg. 1992;47(5):614–20. Epub 1992/11/01. doi: 10.4269/ajtmh.1992.47.614. PubMed PMID: 1449202.

77. Iseki M, Broderson JR, Pirl KG, Igarashi I, Collins WE, Aikawa M. Renal pathology in owl monkeys in Plasmodium falciparum vaccine trials. Am J Trop Med Hyg. 1990;43(2):130–8. Epub 1990/08/01. doi: 10.4269/ajtmh.1990.43.130. PubMed PMID: 2202223.

78. Houba V. Immunopathology of nephropathies associated with malaria. Bull World Health Organ. 1975;52(2):199–207. Epub 1975/01/01. PubMed PMID: 1083308; PubMed Central PMCID: PMCPMC2366359.

79. Chalifoux LV, Bronson RT, Sehgal P, Blake BJ, King NW. Nephritis and hemolytic anemia in owl monkeys (Aotus trivirgatus). Vet Pathol. 1981;18(Suppl 6):23–37. Epub 1981/04/01. doi: 10.1177/0300985881018s0603. PubMed PMID: 7344245.

80. Sato Y, Yanagita M. Renal anemia: from incurable to curable. Am J Physiol Renal Physiol. 2013;305(9):F1239–48. Epub 2013/07/26. doi: 10.1152/ajprenal.00233.2013. PubMed PMID: 23884144.

81. Weller RE. Infectious and Noninfectious Diseases of Owl Monkeys. In: Baer JFW, R.E.; Kakoma, I., editor. Aotus the Owl Monkey. San Diego, CA, USA.: Academic Press, INC.; 1994. p. 178-211

82. Voller A, Davies DR, Hutt MS. Quartan malarial infections in Aotus trivirgatus with special reference to renal pathology. Br J Exp Pathol. 1973;54(5):457–68. Epub 1973/10/01. PubMed PMID: 4202259; PubMed Central PMCID: PMCPMC2072562.

83. Hutt MS, Davies DR, Voller A. Malarial infections in Aotus trivirgatus with special reference to renal pathology. II. P. falciparum and mixed malaria infections. Br J Exp Pathol. 1975;56(5):429–38. Epub 1975/10/01. PubMed PMID: 813757; PubMed Central PMCID: PMCPMC2072788.

84. Silva-Filho JL, Dos-Santos JC, Judice C, Beraldi D, Venugopal K, Lima D, et al. Total parasite biomass but not peripheral parasitaemia is associated with endothelial and haematological perturbations in Plasmodium vivax patients. Elife. 2021;10. Epub 20210929. doi: 10.7554/eLife.71351. PubMed PMID: 34585667; PubMed Central PMCID: PMCPMC8536259.

85. Fonseca JA, Cabrera-Mora M, Singh B, Oliveira-Ferreira J, da Costa Lima-Junior J, Calvo-Calle JM, et al. A chimeric protein-based malaria vaccine candidate induces robust T cell responses against Plasmodium vivax MSP119. Sci Rep. 2016;6:34527. Epub 20161006. doi: 10.1038/srep34527. PubMed PMID: 27708348; PubMed Central PMCID: PMCPMC5052570.

86. Ford A, Kepple D, Abagero BR, Connors J, Pearson R, Auburn S, et al. Whole genome sequencing of Plasmodium vivax isolates reveals frequent sequence and structural polymorphisms in erythrocyte binding genes. PLoS Negl Trop Dis. 2020;14(10):e0008234. Epub 20201012. doi: 10.1371/journal.pntd.0008234. PubMed PMID: 33044985; PubMed Central PMCID: PMCPMC7581005.

87. Kar S, Sinha A. Plasmodium vivax Duffy Binding Protein-Based Vaccine: a Distant Dream. Front Cell Infect Microbiol. 2022;12:916702. Epub 20220713. doi: 10.3389/fcimb.2022.916702. PubMed PMID: 35909975; PubMed Central PMCID: PMCPMC9325973.

88. Jemmely NY, Niang M, Preiser PR. Small variant surface antigens and Plasmodium evasion of immunity. Future Microbiol. 2010;5(4):663–82. doi: 10.2217/fmb.10.21. PubMed PMID: 20353305.

89. Fernandez-Becerra C, Bernabeu M, Castellanos A, Correa BR, Obadia T, Ramirez M, et al. Plasmodium vivax spleen-dependent genes encode antigens associated with cytoadhesion and clinical protection. Proc Natl Acad Sci U S A. 2020;117(23):13056–65. Epub 20200521. doi: 10.1073/pnas.1920596117. PubMed PMID: 32439708; PubMed Central PMCID: PMCPMC7293605.

90. NRC. Guide for the Care and Use of Laboratory Animals. (US) NAP, editor. Washington DC: National Research Council (US) Institute for Laboratory Animal Research; 1996.

91. Ma NS, Rossan RN, Kelley ST, Harper JS, Bedard MT, Jones TC. Banding patterns of the chromosomes of two new karyotypes of the owl monkey, Aotus, captured in Panama. J Med Primatol. 1978;7(3):146–55. Epub 1978/01/01. PubMed PMID: 101668.

92. Obaldia N. Long-term effect of a simple nest-box on the reproductive efficiency and other life traits of an Aotus lemurinus lemurinus monkey colony: an animal model for malaria research. J Med Primatol. 2011. Epub June 18, 2011.

93. Cooper RD. Studies of a chloroquine-resistant strain of Plasmodium vivax from Papua New Guinea in Aotus and Anopheles farauti s.l. J Parasitol. 1994;80(5):789–95. Epub 1994/10/01. PubMed PMID: 7931914.

94. Sullivan JS, Morris CL, Richardson BB, Galland GG, Jennings VM, Kendall J, et al. Adaptation of the AMRU-1 strain of Plasmodium vivax to Aotus and Saimiri monkeys and to four species of anopheline mosquitoes. J Parasitol. 1999;85(4):672–7. Epub 1999/08/26. PubMed PMID: 10461947.

95. Obaldia N, Rossan R, Cooper R, Kyle D, Nuzum E, Rieckmann K, et al. WR 238605, chloroquine, and their combinations as blood schizonticides against a chloroquine-resistant strain of Plasmodium vivax in Aotus monkeys. Am J Trop Med Hyg. 1997;56:508–10.

96. Festing MF, Altman DG. Guidelines for the design and statistical analysis of experiments using laboratory animals. ILAR J. 2002;43(4):244–58. Epub 2002/10/23. doi: 10.1093/ilar.43.4.244. PubMed PMID: 12391400.

97. McCarthy JS, Griffin PM, Sekuloski S, Bright AT, Rockett R, Looke D, et al. Experimentally induced blood-stage Plasmodium vivax infection in healthy volunteers. J Infect Dis. 2013. Epub 2013/08/03. doi: 10.1093/infdis/jit394. PubMed PMID: 23908484.

98. Earle WC, Perez M. Enumeration of parasites in the blood of malarial patients. J Lab Clin Med. 1932:1124–30.

99. Andrysiak PM, Collins WE, Campbell GH. Concentration of Plasmodium ovale- and Plasmodium vivax-infected erythrocytes from nonhuman primate blood using Percoll gradients. Am J Trop Med Hyg. 1986;35(2):251–4. Epub 1986/03/01. PubMed PMID: 3006527.

100. Zhou AE, Berry AA, Bailey JA, Pike A, Dara A, Agrawal S, et al. Antibodies to Peptides in Semiconserved Domains of RIFINs and STEVORs Correlate with Malaria Exposure. mSphere. 2019;4(2). Epub 20190320. doi: 10.1128/mSphere.00097-19. PubMed PMID: 30894432; PubMed Central PMCID: PMCPMC6429043.

101. van den Hoogen LL, Walk J, Oulton T, Reuling IJ, Reiling L, Beeson JG, et al. Antibody Responses to Antigenic Targets of Recent Exposure Are Associated With Low-Density Parasitemia in Controlled Human Plasmodium falciparum Infections. Front Microbiol. 2018;9:3300. Epub 20190116. doi: 10.3389/fmicb.2018.03300. PubMed PMID: 30700984; PubMed Central PMCID: PMCPMC6343524.

102. Oyola SO, Ariani CV, Hamilton WL, Kekre M, Amenga-Etego LN, Ghansah A, et al. Whole genome sequencing of Plasmodium falciparum from dried blood spots using selective whole genome amplification. Malar J. 2016;15(1):597. Epub 20161220. doi: 10.1186/s12936-016-1641-7. PubMed PMID: 27998271; PubMed Central PMCID: PMCPMC5175302.

103. Koren S, Walenz BP, Berlin K, Miller JR, Bergman NH, Phillippy AM. Canu: scalable and accurate long-read assembly via adaptive k-mer weighting and repeat separation. Genome Res. 2017;27(5):722–36. Epub 20170315. doi: 10.1101/gr.215087.116. PubMed PMID: 28298431; PubMed Central PMCID: PMCPMC5411767.

104. Hu G, Llinas M, Li J, Preiser PR, Bozdech Z. Selection of long oligonucleotides for gene expression microarrays using weighted rank-sum strategy. BMC Bioinformatics. 2007;8:350. doi: 10.1186/1471-2105-8-350. PubMed PMID: 17880708; PubMed Central PMCID: PMC2099447.

105. Bozdech Z, Mok S, Gupta AP. DNA microarray-based genome-wide analyses of Plasmodium parasites. Methods Mol Biol. 2013;923:189–211. doi: 10.1007/978-1-62703-026-7_13. PubMed PMID: 22990779.

106. Ritchie ME, Phipson B, Wu D, Hu Y, Law CW, Shi W, et al. limma powers differential expression analyses for RNA-sequencing and microarray studies. Nucleic Acids Res. 2015;43(7):e47. Epub 20150120. doi: 10.1093/nar/gkv007. PubMed PMID: 25605792; PubMed Central PMCID: PMCPMC4402510.

