## Supplementary figures and images for "Sterile protection against *Plasmodium vivax* malaria by repeated blood stage infection in a non-human primate model"

### Figure S1

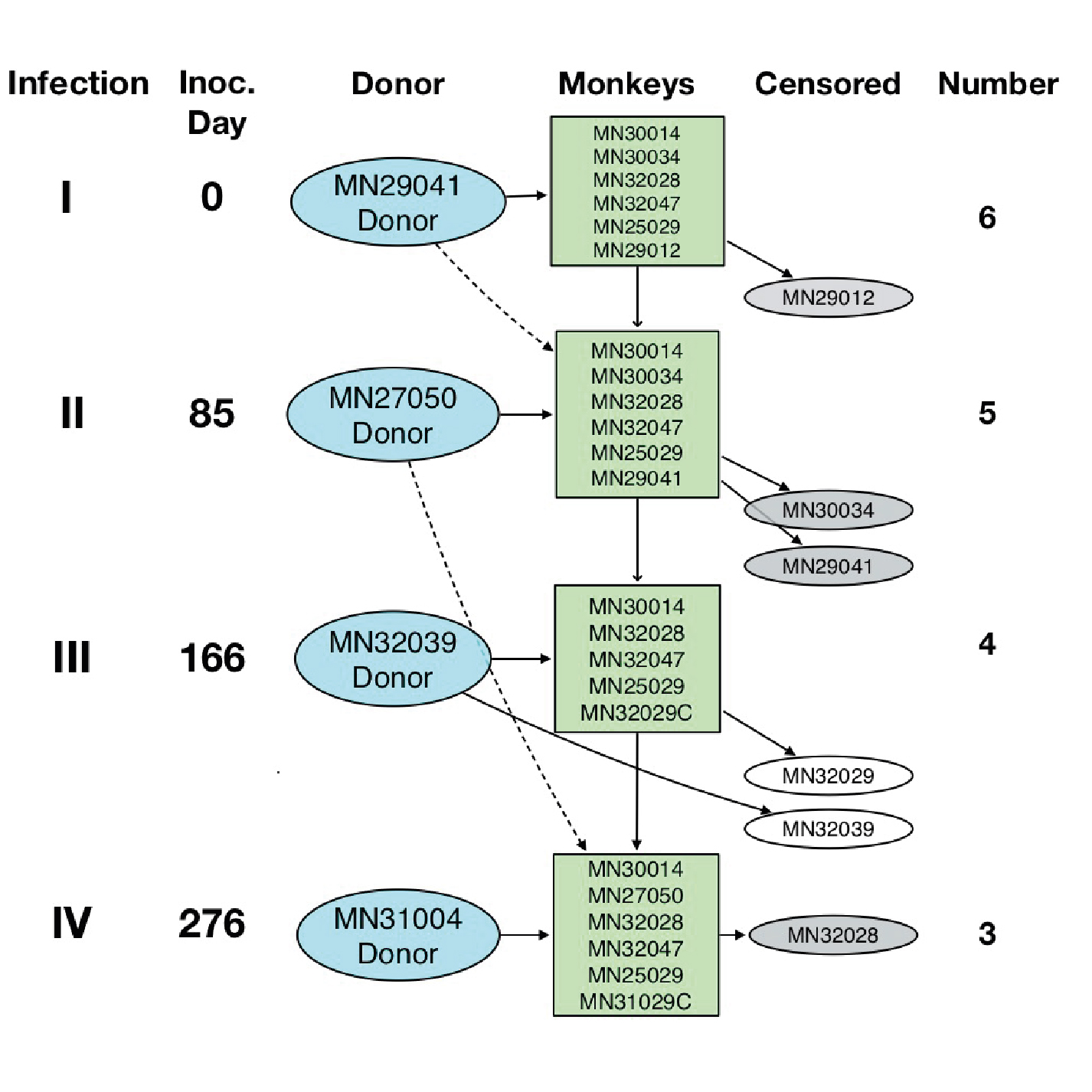

### Figure S2

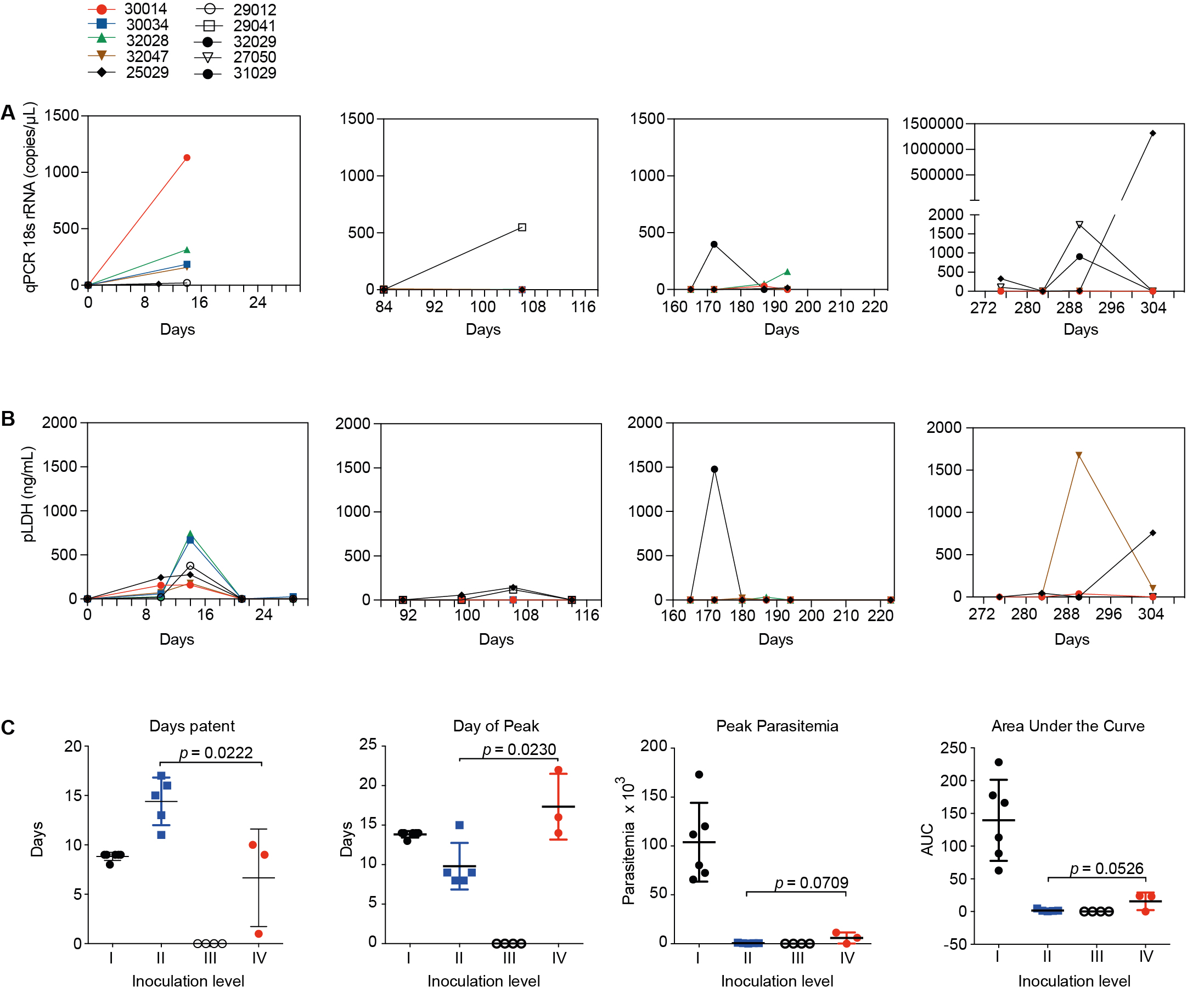

### Figure S3

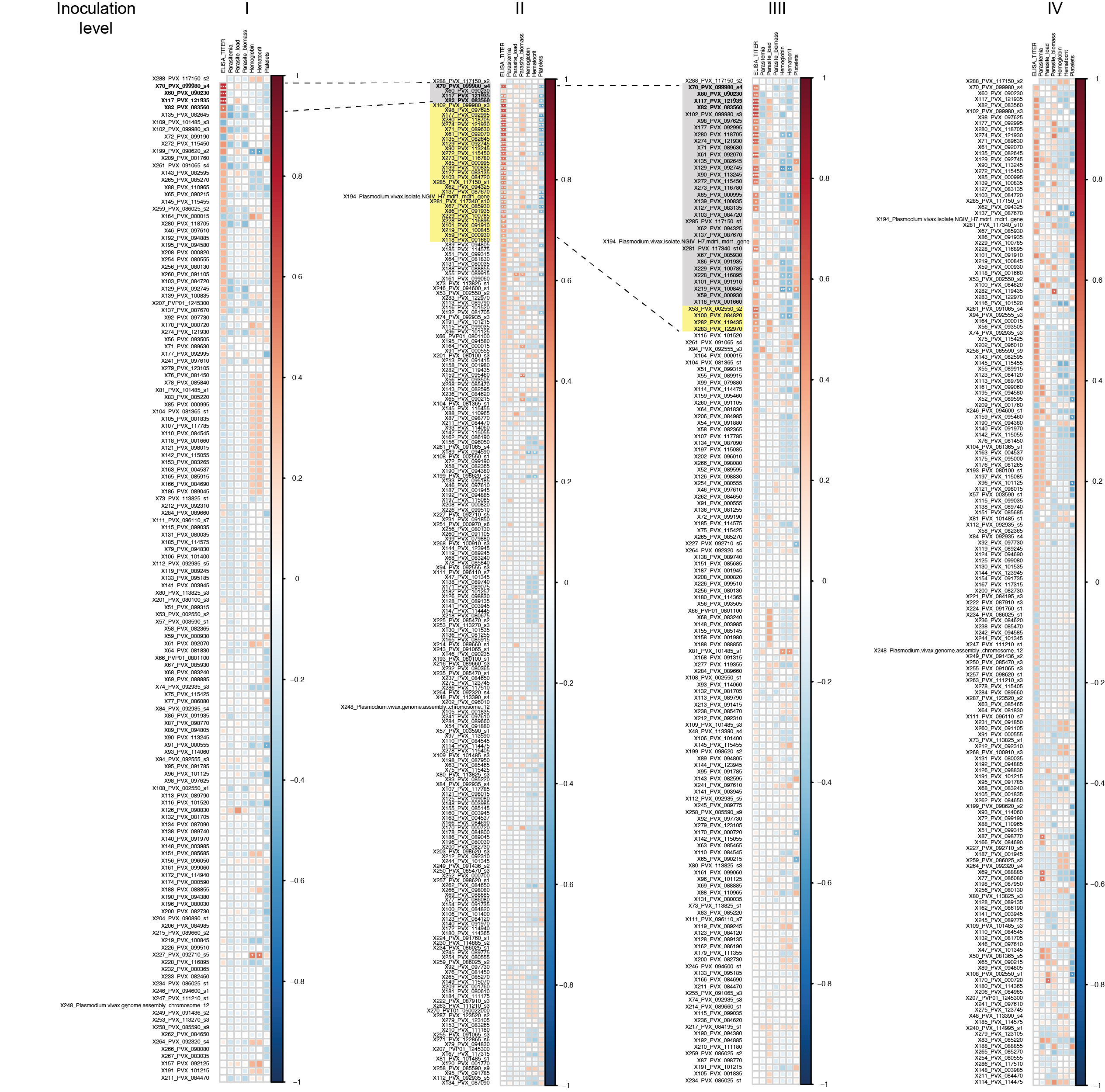

### Figure S4

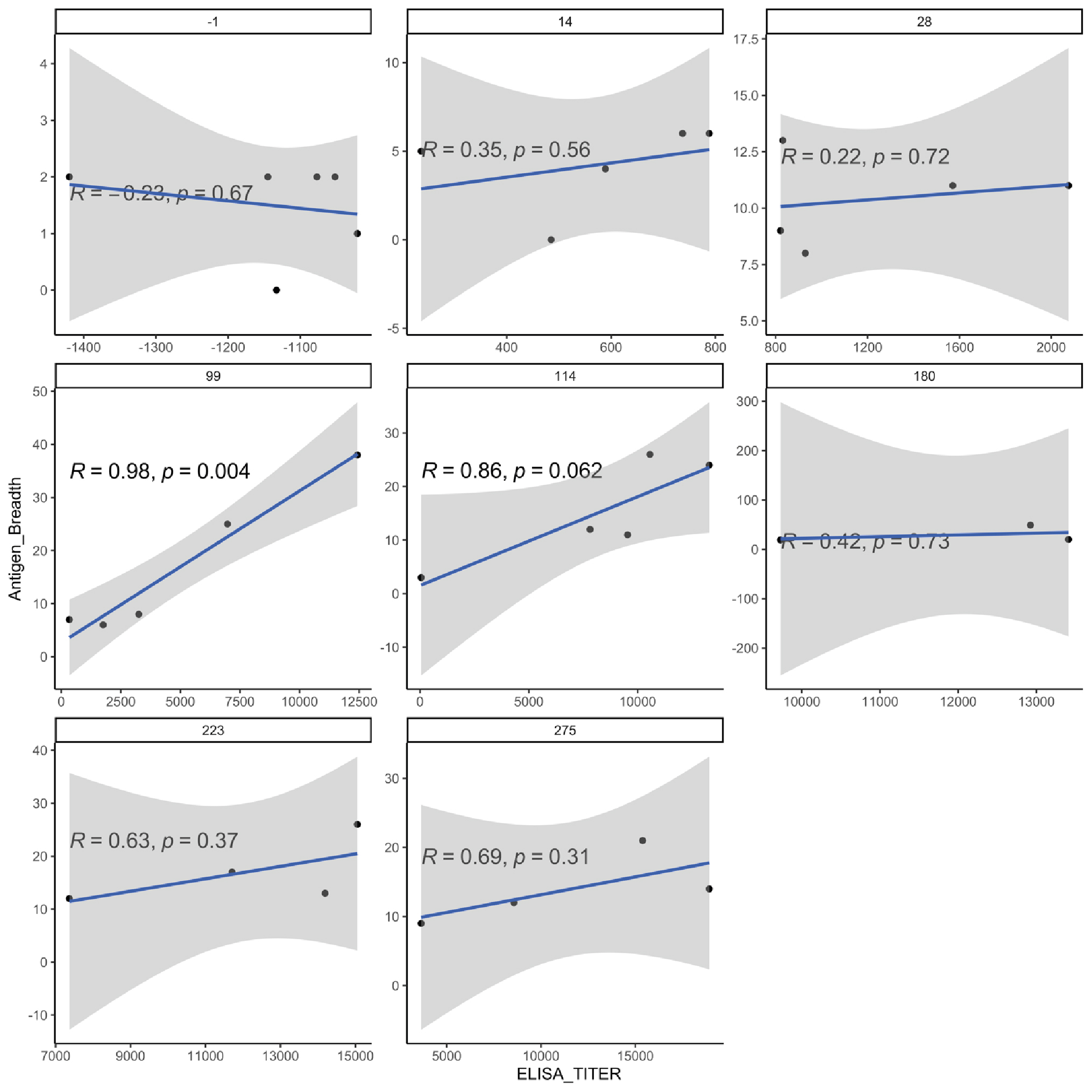

### Figure S5

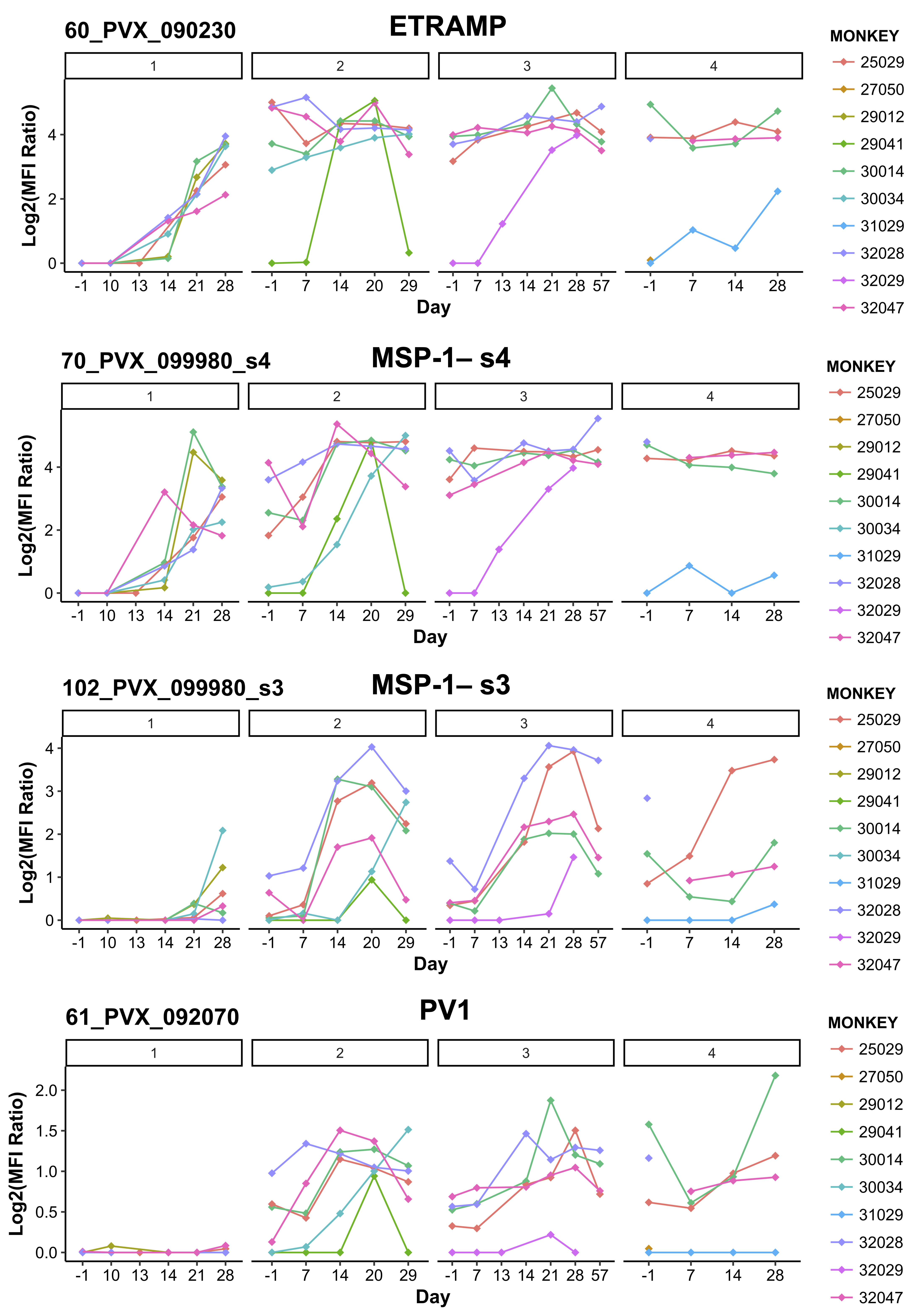
